# Early Host-Virus RNA Interactions Reveal SPEN-Driven m^6^A Regulation as a Major Determinant of Henipavirus Infection

**DOI:** 10.1101/2025.11.21.689838

**Authors:** Andrea Ascura, Stephen Clarke, Andrew F. Livingston, Jackson B. Trotman, Sander Haigh, Morad C. Malek, J. Mauro Calabrese, Manuel Ascano

## Abstract

Early interactions between viral RNA and host-encoded RNA-binding proteins are pivotal in shaping the trajectory of RNA virus infection. Henipaviruses are emerging, highly lethal BSL-4 pathogens whose mechanisms of pathogenesis remain largely elusive. To illuminate the earliest moments of host-virus interplay, we employed Viral Cross-linking and Solid-phase Purification (VIR-CLASP) to capture host proteins bound to the incoming henipavirus genome within the first hour of infection. This approach establishes the first henipavirus RNA-host protein interactome, revealing 146 human proteins directly associated with the primary viral RNA. Among these, SPEN, RBM15, and RBM15B - canonical regulators of lncRNA *Xist* – emerged as key host factors that actively promote viral infection. Direct RNA sequencing further uncovered that SPEN depletion induces widespread hypomethylation, affecting ~98% of differentially modified m^6^A sites, ~87% of which localize to the *L* mRNA transcript encoding the viral RNA-dependent RNA polymerase. Collectively, these findings expose a critical layer of host dependency at the very onset of infection and reveal a previously unappreciated role for SPEN family proteins in facilitating henipavirus infection.

## INTRODUCTION

Emerging infectious diseases continue to pose a profound threat to global public health. Among them, RNA viruses stand out for their remarkable adaptability and role in driving many of the major epidemics and pandemics of the past century. Within this expanding landscape of viral threats, henipaviruses have drawn particular concern, earning classification by the World Health Organization as priority pathogens for research and countermeasure development^1^.

Henipaviruses are zoonotic RNA viruses that include the prototypical BSL-4 agents Hendra virus (HeV) and Nipah virus (NiV)^2,3^. In humans, infection can trigger acute respiratory distress and severe neurologic disease, resulting in recurrent outbreaks across Australia and Asia with alarmingly high case fatality rates^4,5^. The continued discovery of additional henipavirus species across diverse regions, including North America^6^, underscores their global emergence and ecological adaptability^7^. Despite their clinical and epidemiological urgency, no approved therapeutics or vaccines exist for human use^8^, and experimental investigation remains constrained by strict biosafety requirements. Consequently, the molecular mechanisms underlying henipavirus pathogenesis remains largely uncharted. To overcome these barriers, we investigated Cedar virus (CedV) – a henipavirus with high phylogenetic similarity to HeV and NiV - as a tractable BSL-2 model^9,10^ to elucidate the molecular determinants of henipavirus infection and host response.

Henipaviruses are negative-sense, non-segmented RNA viruses that encode for six structural proteins: nucleoprotein (N), matrix protein (M), fusion protein (F), glycoprotein (G), large (L) polymerase protein, and phosphoprotein (P). The viral ribonucleoprotein (RNP) complex consists of the viral RNA genome, N, P, and L protein. Additionally, the P gene encodes additional nonstructural proteins (V/W/C)^11^ (Figure 1A). Henipavirus infection is initiated by the binding of the viral glycoprotein to cellular ephrin B receptors^12–14^ and conformational change of the F protein to induce viral-cell membrane fusion^15^, leading to the release of the viral RNP into the host cytoplasm. The viral polymerase complex, consisting of the P and L protein, first transcribes the encapsidated RNA genome into mRNA then replicates a full-length antigenome, a complementary strand of the viral genome. This antigenome is then used as a template to synthesize copies of the viral genomic RNA^16^.

**Figure 1.**
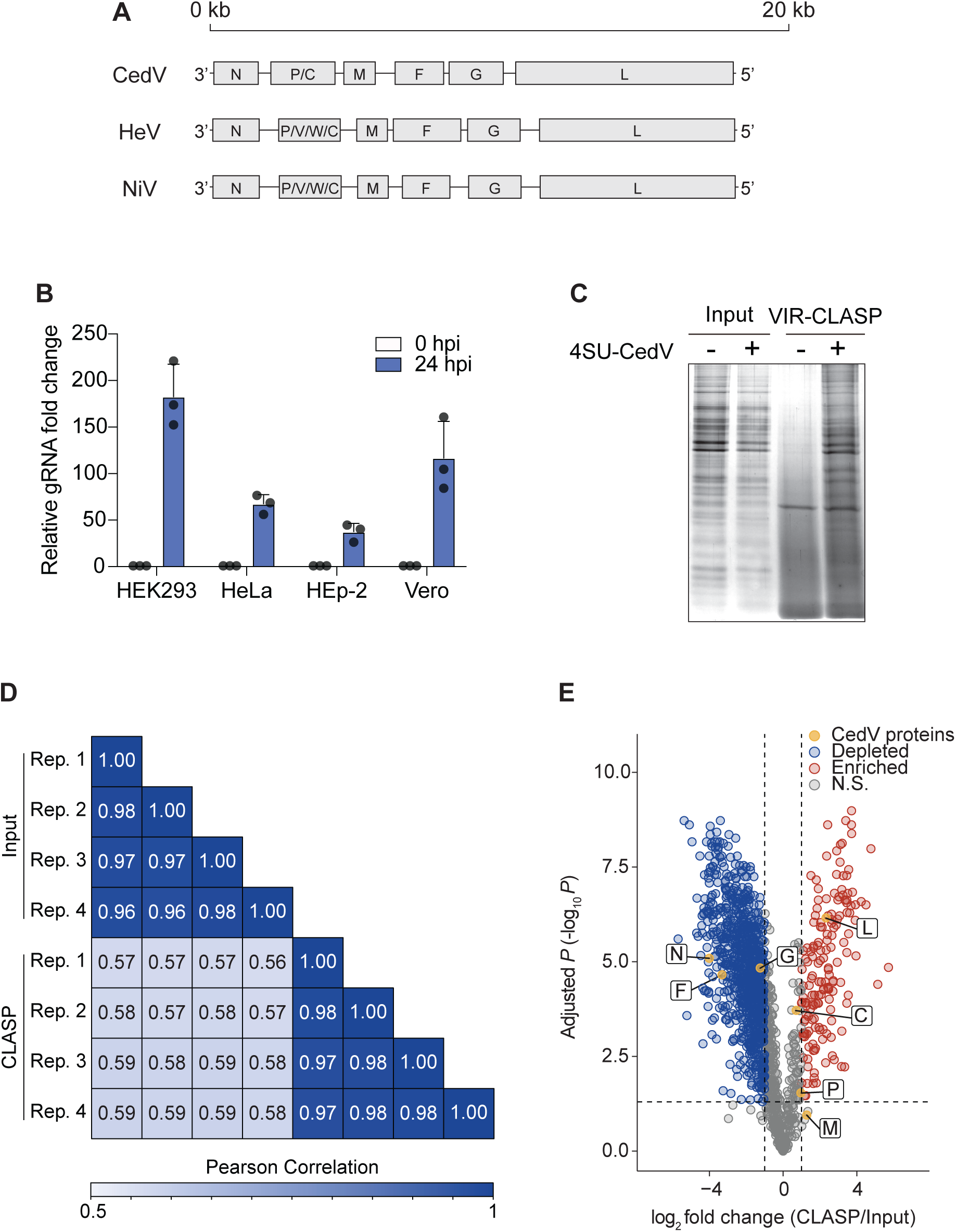
Capturing CedV RNA-protein interactions during pioneer events. (A) Schematic of henipavirus genome for CedV, HeV, and NiV. (B) Strand-specific RT-qPCR of intracellular gRNA from HEK293, HeLa, HEp-2, and Vero cells infected with CedV (MOI 10). Cell lysates were collected at 0- and 24-hours post-infection (hpi). Data is normalized to 0 hpi for each cell line. n = 3, error bars, mean ± SD. (C) SDS-Page and silver stain of proteins purified from HEK293 cells infected with or without 4SU-labeled CedV using VIR-CLASP after 1 hpi. Data is representative of two biological replicates. (D) Heatmap summarizing Pearson correlation coefficients for peptide intensities between Input and CLASP replicates. n = 4, two biological replicates with two technical replicates each. (E) Volcano plot of differentially abundant proteins captured in HEK293 cells infected with 4SU-labeled CedV using VIR-CLASP. Adjusted p values were calculated using student’s t test with Benjamini and Hochberg method and 0.05% false discovery rate cutoff. n = 4.

During infection, henipavirus immune evasion is largely mediated by the P gene nonstructural proteins, V, W, and C. Unlike HeV and NiV which encode all three nonstructural proteins, CedV is known to only encode the C protein^9^, which suppresses expression of inflammatory cytokines^17^. Structural proteins, M and N also antagonize the immune response through interaction with TRIM6^18^ and inhibition of STAT complex formation^19^ during NiV and HeV infection. CedV infection has been shown to differentially upregulate host gene expression of pattern recognition receptors (PRRs), including the cytoplasmic RNA-sensing PRRs retinoic acid inducible gene I and melanoma differentiation-associated protein 5, and other interferon response factors, such as interferon-induced protein with the tetratricopeptide repeats family proteins and 2’5’-oligoadenylate synthetase proteins^20^. While viral immune evasion strategies remain a consistent focus of henipavirus research, very few studies have captured host dependency factors on an unbiased, large-scale basis during henipavirus^21^ or even *Paramyxovirus*^22^ infection. Importantly, no studies have directly interrogated early intracellular events that define henipavirus infection.

Upon entry, incoming viral genomes can be countered by a range of innate immune factors that constitute the host’s intracellular line of defense, preceding the induction of broader antiviral responses^23,24^. During this critical window, viral RNAs (vRNAs) must also engage select host RNA-binding proteins (RBPs) to initiate its own transcription, translation, and replication. Our previous work revealed that the incoming Chikungunya virus (CHIKV) RNA genome recruits numerous host factors including DNA-sensor interferon-inducible protein 16 (IFI16) and N6-methyladenosine (m^6^A) readers YTH domain-containing family (YTHDF) to facilitate CHIKV infection^25^. While we have demonstrated the crucial role of vRNA-host protein interactions during pioneer events, previous studies have highlighted the activity of conventional and unconventional host RBPs in vRNA regulation. These include characterization of SND1, an endonuclease in miRNA decay, as an interactant of negative-sense SARS-CoV-2^26^, GEMIN5, survival motor neuron complex constituent, as a binder of 5’ UTR Sindbis virus transcripts^27^, and ubiquitination of RIG-I by TRIM25 during innate immune sensing^28^. More broadly, viral genomes can employ a range of regulatory events that define cellular processes, including cellular transport^29^, epitranscriptome modulation^30^, and miRNA exploitation^31^. Collectively, these studies highlight the richness of the cellular landscape that can potentially act upon and be exploited by viruses. While the host protein-vRNA interactions has been uncovered for other RNA viruses^25,27,32–34^, cellular interactions with pre-replicative RNA genomes remains an understudied area of vRNA-RBP regulation. Further, the molecular mechanisms by which henipavirus RNA directly interfaces with host factors remain completely uncharacterized.

Here, we employ VIRal-Crosslinking and Solid-phase Purification (VIR-CLASP) to interrogate primary interactions between host proteins and the pre-replicated CedV RNA genome. We discover ~150 human proteins that directly bind CedV. Functional enrichment analyses of the CedV interactome reveal an overrepresentation of various biological categories ranging from RNA regulatory pathways, including splicing and m^6^A methyltransferase complex, to genome maintenance and surveillance. Next, we explored the impact of CedV interacting proteins on infectious virion production and uncovered a subset of candidates that exhibited antiviral or proviral activity.

Notably, we uncovered SPEN family proteins, SPEN (*split ends* gene encoding Msx2-interacting protein, MINT), RBM15, and RBM15B (RNA binding motif protein 15/15B), which are best known for their direct association with long noncoding RNA (lncRNA) *Xist*, as direct vRNA interactants and robust proviral host factors required for viral transcription, replication, and infectious virion production. Given RBM15 and RBM15B’s involvement in the m^6^A RNA methylation of *Xist*, we explored whether CedV infection may also be dependent on m^6^A machinery. We determined that CedV RNA synthesis requires m^6^A methyltransferase writer protein, METTL3, and m^6^A-binding reader proteins, YTHDF1-3. Using nanopore direct RNA-sequencing (DRS), we establish that CedV contains m^6^A modifications across all its mRNA transcripts. Upon depletion of SPEN, m^6^A modifications are dramatically and selectively hypomethylated within the viral L mRNA transcript encoding the RNA-dependent RNA polymerase (RdRp). Overall, our study establishes a molecular framework for dissecting the intracellular mechanisms that govern CedV infection. By identifying pioneer determinants of CedV-host interactions, we not only reveal key regulators of early vRNA metabolism but also uncover a multitude of intrinsic host factors that are poised to deepen our mechanistic understanding of more highly pathogenic henipaviruses. Finally, by defining a role for SPEN family proteins in directing m^6^A deposition on CedV mRNA, our work expands the functional scope of this protein family– long viewed through the lens of *Xist* and X-inactivation - into a previously unrecognized dimension of RNA virus biology.

## RESULTS

### Capturing CedV RNA-protein interactions during pioneer events

To determine a cellular model to investigate pre-replicative interactions during CedV infection, we selected three human cell lines, HEK293, HeLa, and HEp-2 and measured expression of ephrin-B1 (EFNB1) and ephrin-B2 (EFNB2), which are the primary cellular receptors that mediate viral entry^35^. Previous studies have shown that EFNB2-deficient HeLa-USU cells permit CedV entry when transiently expressing EFNB2^9,10^. Additionally, CedV infection of HeLa-CCL2 cells has been shown to induce an immune response 24 hours post-infection (hpi)^20^. Nevertheless, in addition to HeLa cells, we selected additional human lines, HEK293 and HEp-2, to determine whether human cell lines used for RNA virus infection might also prove susceptible and permissive to CedV infection^27,36^. We also included African green monkey cell line, Vero cells, which is an established line used for CedV propagation and amplification^9,10^, as an additional positive control. In uninfected conditions, we observe that both HEK293 and Vero cells expressed higher expression of both EFNB1 and EFNB2 when compared to that of HeLa cells. Specifically, HEK293 and Vero showed the greatest expression of EFNB1 and EFNB2 by RT-qPCR, respectively (Figure S1A). We next determined whether these cell lines supported viral replication by measuring CedV genomic RNA (gRNA) following 24 hours of infection. Although all tested cell lines permitted CedV replication, infection in HEK293 cells resulted in the most robust viral replication; ~4-fold greater compared to that of HeLa cells, the first line used for initial CedV studies (Figure 1B). Given these results, we selected HEK293 cells to investigate host-virus interactions during CedV infection.

To profile pioneer interactions between the incoming, pre-replicated genome of CedV and cellular proteins, we performed VIRal-Crosslinking and Solid-phase Purification (VIR-CLASP) on 4-thiouridine (4SU)-labeled CedV^37^. Briefly, CedV was propagated in Vero cells in the presence of 4SU and labeled viral particles were purified by ultra-centrifugation. 4SU incorporation was confirmed by high-performance liquid chromatography and measured to incorporate at 0.8%, or 0.8 4SU for every 100 uridines within the CedV genome (data not shown). Our previous work using 4SU in cellular and viral contexts indicate that this level of 4SU incorporation is sufficient to capture RNP complexes^25,37^. HEK293 cells were infected with 4SU-labeled CedV and irradiated with UV light at 1 hpi to covalently crosslink vRNA and directly bound proteins. By silver-stain, we observed diverse bands in the 4SU-CedV VIR-CLASP sample when compared to the 4SU-CedV Input sample indicating capture of distinct interactions between 4SU-labeled CedV and cellular proteins (Figure 1C).

Next, we subjected VIR-CLASP enriched proteins to liquid chromatography coupled with tandem mass spectrometry (LC-MS/MS) using a label-free quantitation (LFQ) approach, followed by assessment of quality control metrics and proteomic analysis. Enrichment of VIR-CLASP RBPs were calculated by computing protein intensity ratios between proteins found in +4SU CLASP versus +4SU Input samples. As expected, replicates within the same sample group were strongly correlated (Figure 1D). By contrast, comparisons between Input and CLASP replicates were only moderately correlated at the peptide intensity level (Figure 1D). Further, Input and CLASP samples demonstrated distinct proteome compositions despite similar compositions between replicates of the same sample group (Figure S1B-C). Altogether, we determined that our proteomic samples yielded high quality metrics, representing sufficient protein dynamic range and complexity within and across Input and CLASP samples as well as high correlation between matched replicates.

We discovered 1215 distinct human proteins that were statistically significant in our CLASP sample (adjusted *P* < 0.05) (Figure 1E and Table S1). Among these, we identified 1016 proteins that were depleted (log_2_ fold change < 0) and 199 proteins that were enriched (log_2_ fold change > 0). To filter for highly enriched candidates for subsequent analyses, we establish a log_2_ fold change > 1 as our threshold cutoff. For this study, we define a subset of 146 human proteins as the CedV RNA interactome (CedV VIR-CLASP proteins).

Approximately 72% (107/147 proteins) of our interactome has been previously identified in other vRNP-host protein studies, thus demonstrating the specificity of our crosslinking methodology, VIR-CLASP, to capture vRNA interactants. Nevertheless, we also observe that ~28% (41/147 proteins) of our interactome represent putative cellular RBPs that are potentially unique interactants of incoming CedV RNA and uncharacterized regulators of vRNA (Figure S1D and Table S2).

In addition to enriched human VIR-CLASP proteins, we identified 3 of 7 CedV proteins – L, C, and P - that were enriched (log_2_ fold change > 0) (Figure 1E). The L and P protein comprise the CedV RdRp machinery, which is essential for transcribing and replicating vRNA^38,39^. We found that in addition to capturing the P protein, the L protein was the most enriched among viral proteins identified. Although the nonstructural C protein has numerous previously reported activities during henipavirus pathogenesis, including modulation of host proinflammatory response and promotion of virus budding, the exact role of C in directly binding vRNA during infection remains unclear^17,40^. Among all the viral proteins, the N protein was the most depleted in our dataset. This was a key finding since the N protein is known to encase the viral genome. Its depletion is consistent with numerous structural studies that propose a model of nucleocapsid uncoiling and displacement to allow for viral polymerase accessibility and progression^41–43^. Thus, our results appear to show that we have captured a pivotal timepoint in which the viral RdRp, the most enriched viral protein in our dataset, is engaged on the incoming genome alongside intracellular host factors.

### Functional enrichment analysis of the pre-replicated CedV interactome

We performed functional enrichment analyses using the CedV interactome to understand the composition of proteins interacting with the incoming CedV RNA genome. Pathway enrichment analysis using the Reactome database revealed an overrepresentation of cellular and viral pathways spanning transcription-and translation-related processes^44^ (Figure S2A-B and Table S3). The enrichment of both cellular- and virus-dependent processes underscores the central paradigm that vRNA must appropriate host regulatory pathways to drive infection^25,45,46^. Notably, host proteins in the interactome enrich for pathways annotated for other viruses, including SARS-CoV-2, HIV, and Influenza (Figure S2B).

Next, we further explored the connectivity of RBPs within our interactome to determine whether there was an enrichment of functionally important clusters. We constructed a protein-protein association network and evaluated the enrichment of GO terms within each cluster^47,48^. Overall, annotations within the network can be broadly categorized as RNA metabolism, gene expression, and genome maintenance and surveillance (Figure 2A, Figure S2C, and Table S3). We observe a strong enrichment of terms associated with RNA metabolism, such as splicing^49,50^, mRNA transport^29,51,52^, and m^6^A methyltransferase complex^53–55^, which have been studied in other RNA viruses but have yet to be well characterized for henipaviruses.

**Figure 2.**
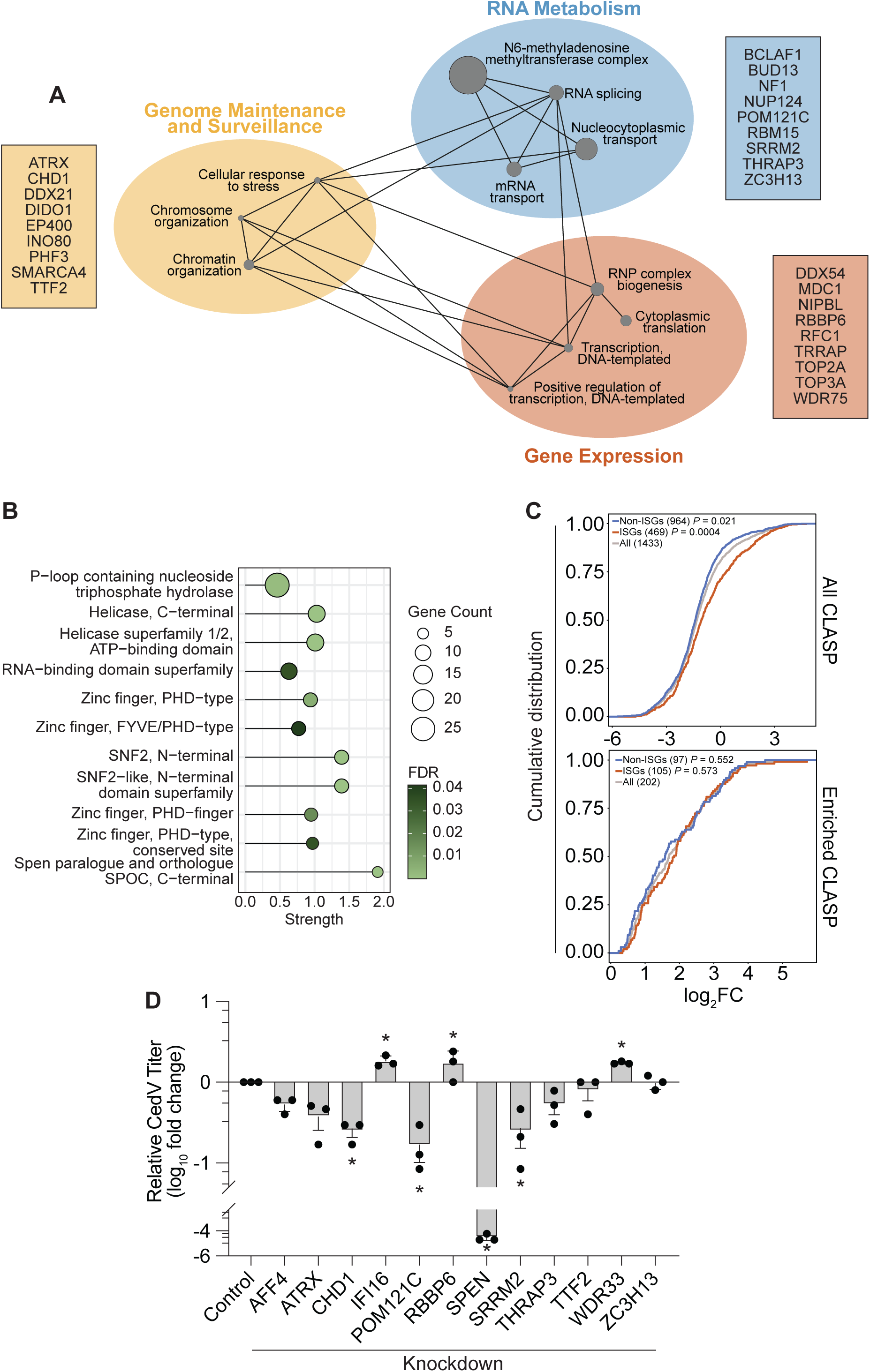
Functional enrichment analysis of the pre-replicated CedV interactome. (A) Protein-protein association network of CedV RNA interactome retrieved from STRING v12.0^48^. Each grey node represents a specific GO term and is scaled to the strength, defined as log_10_(observed/expected), of the enrichment effect. Enrichment effect strength is calculated based on the number of proteins in the network with the annotated term relative to the number of proteins expected to be annotated with the given term in a random network of the same size. Colored circles represent closely related GO terms. Connections represent shared genes. Boxes display a subset of candidates in the shared functional category. (B) Dot plot of enriched InterPro domains and features in the CedV RNA interactome. Enrichment is based on all human proteins. Circles are scaled to gene count for each protein feature. Terms are sorted by gene count. FDR is computed with Fishers exact test using Benjamini-Hochberg correction. (C) Top: Cumulative distribution fraction analyses for all CLASP proteins on based on ISG annotations retrieved from Interferome database^68^. Adjusted *P* values were calculated using pairwise Wilcoxon test with Benjamini-Hochberg method. Bottom: Cumulative distribution fraction analyses for enriched CLASP proteins based on ISG annotations. Adjusted *P* values were calculated using pairwise Wilcoxon test with Benjamini-Hochberg method. (D) siRNA knockdown of CedV RNA interactome proteins in HEK293 cells followed by measurement of viral titer. siRNA transfection was performed for 72 h followed by CedV infection (MOI 0.1) for 16 h. Titer was determined by plaque assay in Vero cells. Data is normalized to non-template control. n = 3, error bars, mean ± SD. P values were calculated using one-way Anova with Dunnett’s multiple comparisons test.

We then evaluated the overrepresentation of protein domains and features enriched among CedV VIR-CLASP protens. Using InterPro classifications, we observe an enrichment of proteins domains, such as P-loop containing nucleoside triphosphate hydrolase and helicase superafmily, characterized in nucleotide-binding to either RNA or DNA (Figure 2B and Table S3). These DNA- and RNA-binding proteins, such as IFI16^25,56,57^ and DHX29^58^, are known to control transcription and translation, maintain genome integrity, and mediate cellular stress responses^59,60^. As expected, we also observed an enrichment of the RNA-binding domain superfamily, which includes the RNA recognition motif (RRM) domain. Proteins within this classification, such as DDX21^61^ and HNRNPF^62^, which have been studied in their canonical roles in RNA metabolism. Interestingly, we also found an enrichment of proteins harboring the Spen paralogue and orthologue SPOC domain, such as SPEN, a lncRNA RBP involved in transcriptional regulation^63–66^.

Since there are interferon-stimulated genes (ISGs) RBPs that can exert post-transcriptional regulation during an immune response^67^, we evaluated whether there was an ISG enrichment within our interactome^68^. We examined the cumulative distribution of all 1433 proteins identified in our CLASP sample to discern the probability of ISGs being enriched in our dataset. Our analysis revealed that ISGs were significantly enriched, while non-ISGs were significantly depleted (Figure 2C). When we exclusively examined the distribution of ISGs among enriched CLASP proteins (202 proteins), there was no significant enrichment or depletion of proteins with either ISG or non-ISG annotations. Further, there was a broadly even distribution in the number of proteins annotated in both categories, in which ~52% (105/202 proteins) were annotated as ISGs, while ~48% (97/202 proteins) were annotated as non-ISGs (Figure 2C). These results suggest that the incoming CedV genome does not appear to preferentially recruit ISG RBPs categorically during pioneer interactions of infection.

### VIR-CLASP candidates are important for CedV infectious virion production

We were interested in exploring how a subset of the CedV interactome perturbed various stages of the viral lifecycle. To select CedV RBPs for functional assessment, we leveraged functional enrichment analyses and selected candidates based on a number of criteria. We prioritized strongly enriched proteins that had previous identification as an RBP in either viral or cellular RNP complexes and was implicated in either viral infections or innate immunity. Additionally, the candidates were selected from a broad coverage and representation of the biological categories present within the CedV interactome. Using this criteria, we performed siRNA-induced knockdown on a subset of our interactome that included AFF4, ATRX, CHD1, IFI16, POM121C, RBBP6, SPEN, SRRM2, THRAP3, TTF2, WDR33, and ZC3H13 and evaluated nascent virion production following viral infection (Figure 2D and Figure S3A). Among the candidates tested, CHD1, IFI16, POM121C, RBBP6, SPEN, SRRM2, and WDR33 resulted in a significant change in titer compared to the control. Specifically, IFI16, RBBP6, and WDR33 increased infectious virion production and exhibited antiviral activity against CedV infection (Figure 2D). IFI16, an innate immune sensor for intracellular DNA^56,69^, was previously identified to directly bind CHIKV and Influenza A (IAV) RNA genomes and act as antiviral factors for both viruses^25,57^. WDR33, a component of the 3’ mRNA cleavage and polyadenylation specificity factor complex^70^, was also reported as a direct interactant of CHIKV and IAV pre-replicated genomes^25^. In contrast, we found that CHD1, POM121C, SPEN, and SRRM2 acted as proviral factors that reduced production of infectious virions upon their depletion (Figure 2D). CHD1, a chromatin assembly factor involved in nucleosome spacing^71,72^, was not only identified as a direct binder of IAV genome^25^, but also as an interactant with the IAV polymerase complex that reduced vRNA replication and mRNA transcription upon silencing^73^. POM121C is a structural constituent of the nuclear pore complex^74^ that was shown to inhibit HIV-1 replication^75^. SRRM2, a component of the spliceosome, is involved in pre-mRNA splicing and nuclear speckle formation^76–79^. SRRM2 was also previously identified to bind pre-replicated CHIKV genomes.^25^ Overall, our results expand on these findings by suggesting an unexplored role of these RNA interactants in mediating CedV infection through vRNA binding.

While we uncovered numerous host proteins that exhibited antiviral and proviral activities upon their loss, the most striking impact we observed was that from the depletion of SPEN. Loss of SPEN reduced viral titer by ~100,000-fold and was the most robust host factor among the VIR-CLASP candidates we tested. Given this result and for this report, we focused on characterizing how SPEN, a lncRNA RBP involved in transcriptional silencing^80^, affects the CedV lifecycle.

### SPEN is a robust proviral factor necessary for viral infection

SPEN, also known as MINT or SHARP (<u>S</u>MRT/<u>H</u>DAC1-<u>a</u>ssociated repressor <u>p</u>rotein), was first identified as a recessive lethal mutation in *Drosophila* embryos^81^. Although initially characterized as a transcriptional regulator of neuronal cell fate^82–84^, the biological role of SPEN is largely focused on its epigenetic regulation of the mammalian X-chromosome. SPEN is a canonical RBP that harbors four RNA recognition motifs at the N-terminus and a SPOC domain (<u>S</u>pen <u>p</u>aralog and <u>o</u>rtholog <u>C</u>-terminal domain) at the C-terminus^85,86^. Further, SPEN has been experimentally captured as an RBP using various biochemical methodologies probing RNA-protein interactions, including RIC^87–90^, RBDmap^91^, serIC^92^, pCLAP^93^, eRIC^94^, RICK^95^, OOPS^96^, PTex^97^, and XRANAX^98^. Notably, SPEN was identified as an RNA-binding partner of *Xist*, an essential lncRNA for X chromosome inactivation^66^ and has been further characterized in *Xist* regulation and gene silencing^63–65,99–102^. Intriguingly, SPEN has also been identified in association with retroviral RNA elements^103^ and with other vRNA interactomes^26,104^ – suggesting that its engagement with CedV RNA may reflect a broader, intrinsic RNA-recognition function rather than a henipavirus-specific hijacking mechanism.

We first evaluated whether SPEN was important during earlier stages of the CedV life cycle by utilizing a CedV reporter containing GFP (CedV-GFP) to monitor intracellular viral replication^10^. We infected HEK293 cells with CedV-GFP and monitored fluorescent expression in SPEN knockdown or control conditions using flow cytometry. Loss of SPEN showed a dramatic reduction of GFP protein expression from ~73% in the control to ~5% in SPEN-depleted samples (Figure 3A) and significantly decreased mean GFP intensity (Figure 3B). These results are independent of any effect to cell viability following SPEN depletion (data not shown). Thus, SPEN is important for not only the production of viral progeny, but also for intracellular viral replication.

**Figure 3.**
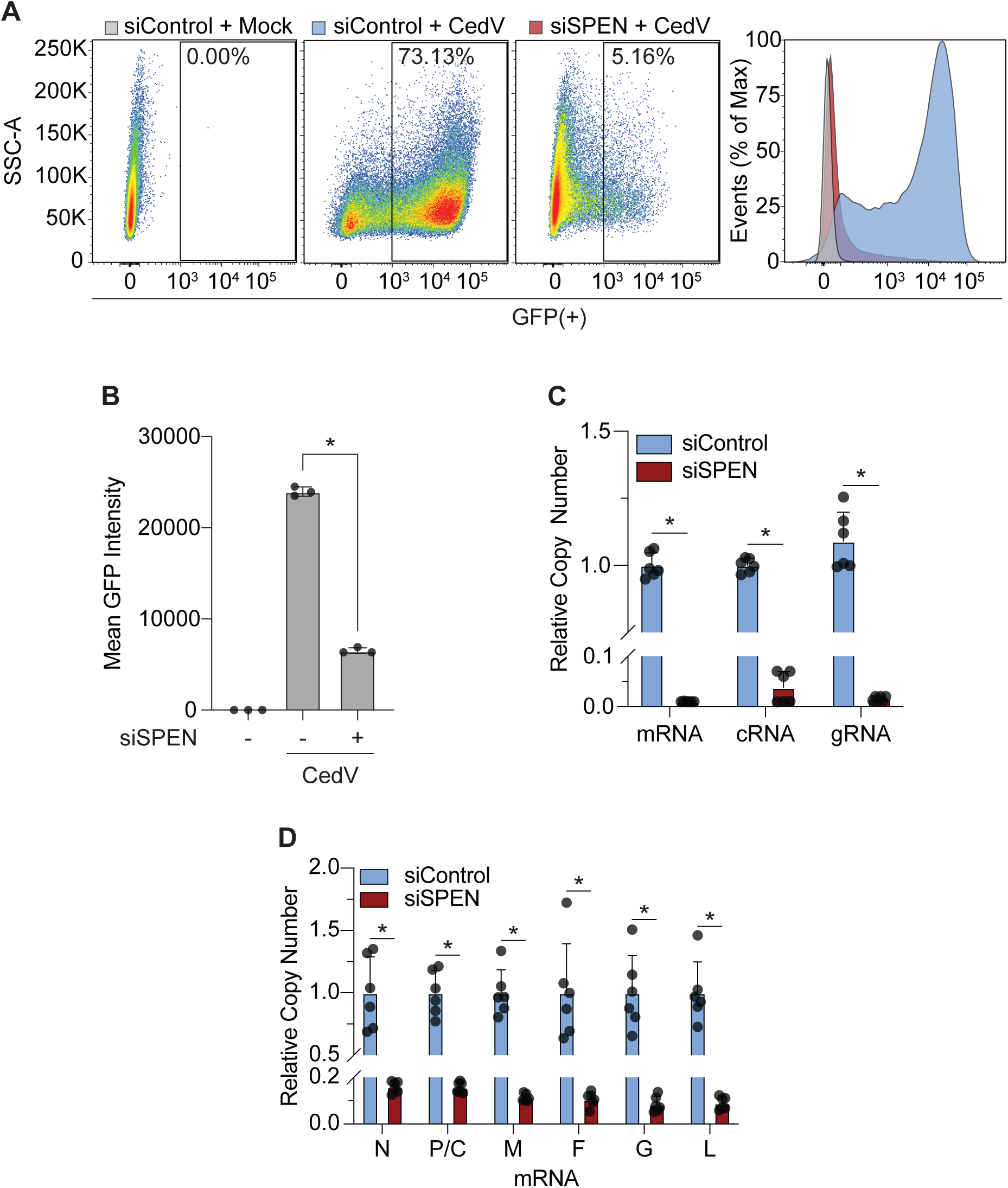
SPEN is a robust proviral factor necessary for viral infection. (A) Left: Scatter plots of GFP expression in SPEN knockdown or control HEK293 cells infected or uninfected with CedV-GFP (MOI 0.1). siRNA transfection was performed for 72 h followed by infection for 16 h. Percentage represents GFP expression gated based on uninfected, control conditions (siControl + Mock). Right: Representative histogram overlays of GFP expression between conditions. n = 3. (B) Mean GFP intensity in SPEN knockdown or control HEK293 infected or uninfected with CedV-GFP (MOI 0.1). n = 3, error bars, mean ± SD. P values were calculated using one-way Anova with Dunnett’s multiple comparisons test. (C) siRNA knockdown of SPEN or control HEK293 cells followed by strand-specific RT-qPCR to measure viral RNA strands. siRNA transfection was performed for 72 h followed by CedV infection (MOI 0.1) for 16 h. Data is normalized to non-template control. n = 6, error bars, mean ± SD. P values were calculated using ordinary two-way Anova with Sidak’s multiple comparisons test. (D) siRNA knockdown of SPEN or control in HEK293 cells followed by strand-specific RT-qPCR to measure CedV mRNA transcripts. siRNA transfection was performed for 72 h followed by CedV infection (MOI 0.1) for 16 h. Data is normalized to non-template control. n = 6, error bars, mean ± SD. P values were calculated using ordinary one-way Anova with Sidak’s multiple comparisons test.

Since loss of SPEN reduced intracellular viral replication, we next determined whether SPEN was required for the synthesis of the three distinct vRNA strands produced during viral transcription and replication. As a negative sense RNA virus, CedV must transcribe its incoming genomic RNA (gRNA) into mRNA using its RdRp. Following transcription, the gRNA is also used to synthesize positive-sense complementary RNA (cRNA), which then serves as the template for replication and amplification of gRNA for viral progeny^16^. We captured these three distinct viral strands (gRNA, mRNA, and cRNA) in control and SPEN knockdown HEK293 cells using strand specific qPCR (Figure S3B-D). SPEN depletion significantly reduced viral mRNA, cRNA, and gRNA and thus restricted viral transcription and replication (Figure 3C). While we evaluated P mRNA as a proxy for strand specific qPCR, we similarly observed that SPEN depletion reduced the expression of other CedV mRNA transcripts (Figure 3D). Overall, our results show that SPEN is a robust proviral factor required for various stages of the viral lifecycle from viral transcription to virion production.

### Additional SPEN family proteins, RBM15 and RBM15B promote viral infection

In addition to identifying SPEN as a vRNA binder, we captured two additional SPEN family protein members, RBM15 and RBM15B, as enriched VIR-CLASP proteins. Owing to their classification as SPEN family proteins, RBM15 and RBM15B harbor identical protein domain organization, consisting of four N-terminal RRMs and a C-terminal SPOC domain^105^. Although RBM15 and RBM15B have also been captured in numerous vRNP complexes^25,26,33,104^, their roles in mediating viral infection have yet to be characterized.

We first determined the functional role of RBM15 and RBM15B in CedV pathogenesis by measuring nascent virion production from CedV-infected siRBM15 and siRBM15B HEK293. Loss of both RBM15 and RBM15B significantly reduced viral titer. Specifically, RBM15B depletion reduced viral titer by ~1,000-fold, while RBM15 depletion resulted in a ~10-fold reduction (Figure 4A). While loss of either RBM15 and RBM15B did not restrict infectious virion production as dramatically as SPEN depletion, both RBM15 and RBM15 are robust proviral host factors when compared to other VIR-CLASP candidates tested in our initial siRNA screen (Figure 2D).

**Figure 4.**
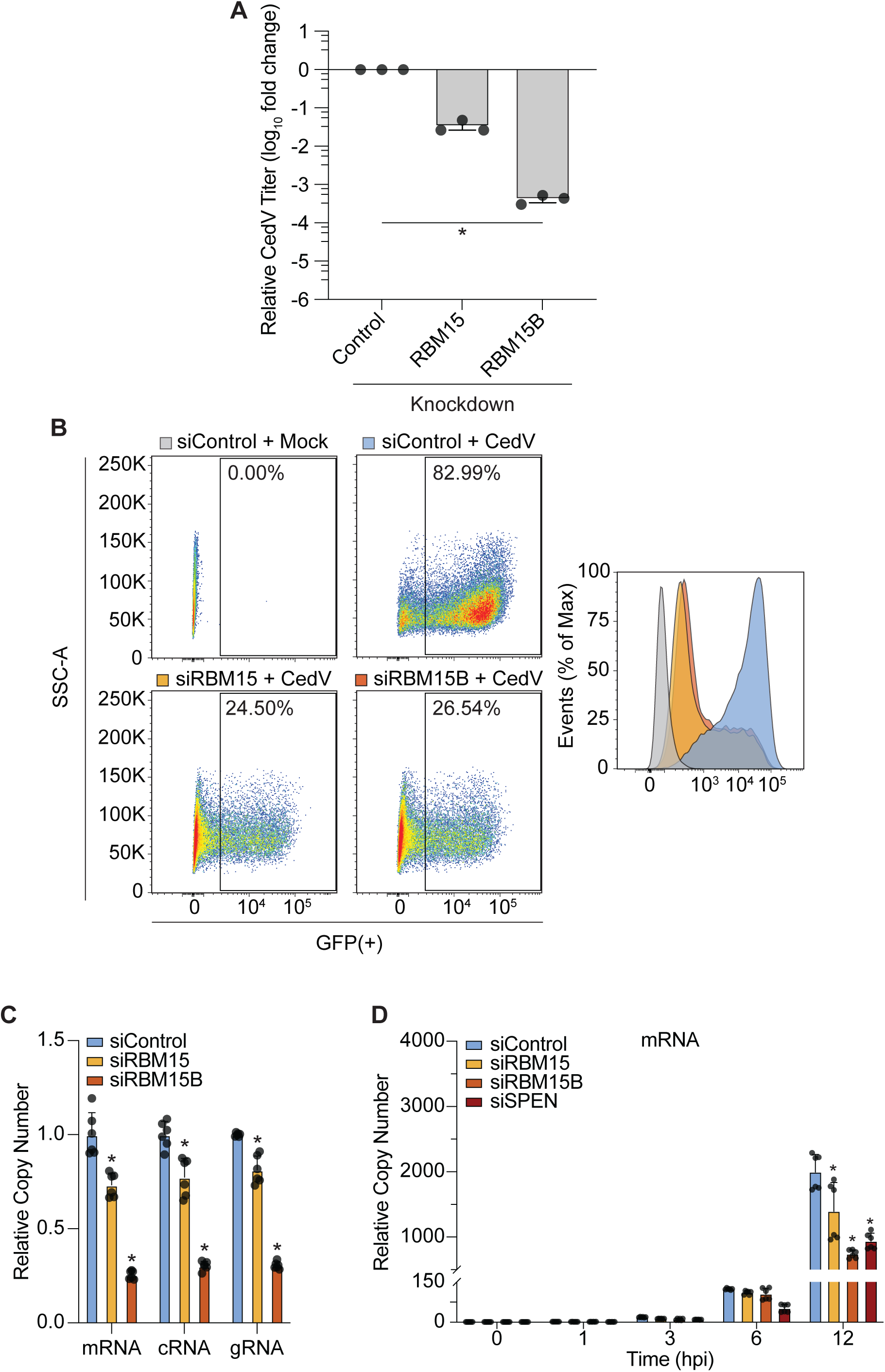
Additional SPEN family proteins, RBM15 and RBM15B promote viral infection. (A) siRNA knockdown of RBM15, RBM15B, or control in HEK293 cells followed by measurement of CedV titer. siRNA transfection was performed for 72 h followed by CedV infection (MOI 0.1) for 16 h. Titer was determined by plaque assay in Vero cells. Data is normalized to non-template control. n = 3, error bars, mean ± SD. P values were calculated using one-way Anova with Dunnett’s multiple comparisons test. (B) Left: Scatter plots of GFP expression in control, RBM15, or RBM15B knockdown HEK293 cells infected or uninfected with CedV-GFP (MOI 0.1). siRNA transfection was performed for 72 h followed by infection for 16 h. Percentage represents GFP expression gated based on uninfected, control conditions. Right: Representative histogram overlays of GFP expression between conditions. n = 3. (C) siRNA knockdown of RBM15, RBM15B, or control in HEK293 cells followed by strand-specific RT-qPCR to measure viral RNA strands. siRNA transfection was performed for 72 h followed by CedV infection (MOI 0.1) for 16 h. Data is normalized to non-template control. n = 6, error bars, mean ± SD. P values were calculated using ordinary two-way Anova with Dunnett’s multiple comparisons test. (D) siRNA knockdown of RBM15, RBM15B, SPEN, and control in HEK293 cells followed by strand-specific RT-qPCR to measure viral mRNA. siRNA transfection was performed for 72 h followed by CedV infection (MOI 0.1) for indicated timepoints. Data is normalized to 0 hpi for each knockdown condition. n = 6, error bars, mean ± SD. P values were calculated using two-way Anova with Dunnett’s multiple comparisons test.

Next, we investigated whether RBM15 and RBM15B were required for intracellular viral replication by using CedV-GFP to monitor fluorescent expression during infection of either siRBM15 or siRBM15B HEK293. We observed that loss of RBM15 or RBM15B dramatically reduced GFP expression by over 50% in either condition. Specifically, RBM15 and RBM15B depletion showed 24.50% and 26.54% GFP expression, respectively, compared to control with ~83% GFP expression (Figure 4B). However, loss of either RBM15 or RBM15B did not reduce GFP expression as dramatically as SPEN depletion alone (Figure 3A). Further, reduction of GFP expression was still greater following loss of SPEN alone compared to simultaneous depletion of both RBM15 and RBM15B (Figure S4B). We observed the most pronounced reduction in GFP intensity following loss of both RBM15 and SPEN or RBM15B and SPEN or loss of all three SPEN family proteins (Figure S4B). Individually, SPEN remains the most robust proviral factor among SPEN family proteins.

We then examined whether RBM15 and RBM15B modulated CedV RNA synthesis by using strand specific qPCR to measure CedV mRNA, cRNA, and gRNA following RBM15 or RBM15B knockdown. Loss of RBM15 and RBM15B significantly reduced levels of CedV gRNA, mRNA, and cRNA (FIG 4C). Using this same approach, we investigated whether SPEN, RBM15, and RBM15B were required for viral transcription and replication during early timepoints of infection. Significant reduction of viral gRNA, mRNA, and cRNA levels following knockdown of either SPEN, RBM15, and RBM15B was observed as early as 12 hpi (Figure 4D and Figure S4D-E).

While our observations following depletion of SPEN, RBM15, and RBM15B are primarily intracellular, we recognize that our results may be due to other consequences, particularly viral entry, caused by loss of SPEN family proteins that might influence subsequent stages of viral infection. Previous genome-wide screens identifying host factors required for viral infection observed that loss of candidate proteins perturbed various stages of the viral lifecycle, ranging from entry to post-entry. Specifically, loss of antiviral factor, USP47, and proviral factor, DPAGT1 was determined to perturb viral entry for IAV^106^ and SARS-CoV-2^107^, respectively, while other candidates were shown to impact post-entry events. To address the possibility that our post-entry observations were due to a perturbation with initial viral entry, we tested EFNB1 and EFNB2 expression following depletion of SPEN, RBM15, and RBM15B. While EFNB2 expression was not significantly altered, EFNB1 expression was increased following depletion of RBM15 (Figure S4C). Although loss of RBM15 led to elevated EFNB1 expression, CedV vRNA levels during early infection were unchanged. This indicates that the modest rise in EFNB1 mRNA does not confer a functional increase in CedV entry (Figure 4D and Figure S4D-E). Additionally, to rule out the possibility that our observations were due to cytotoxicity following SPEN family protein depletion, we evaluated the proportion of live cells across SPEN, RBM15, and RBM15B knockdown conditions using flow cytometry and observed no differences in the percentage of live cells when compared to the control (data not shown). These results confirm our findings that the proviral effects exerted by SPEN, RBM15, and RBM15B are due to intracellular, post-entry activity. Taken together, our results thus far not only demonstrate that RBM15, RBM15B, and SPEN are important proviral factors for overall viral replication but also suggest that there is a potential epistatic relationship underlying the recruitment of these proteins to viral RNA to elicit their function, which merits future investigation.

### Cedar virus transcription and replication is dependent on m^6^A machinery

m^6^A is an abundant RNA modification found in eukaryotic mRNA and viral genomes^108–113^, including NiV RNA from various strains^114^. Methylation of the N6-position of adenosine is deposited by methyltransferases (“writers”), removed by demethylases (“erasers”), and influences the association of a growing list of m^6^A-binding proteins (“readers”)^108^. m^6^A methylation is mediated by methyltransferase complex members, METTL3 and METTL14^115^. Since the initial identification of the first methyltransferase, METTL3, the growing body of research in m^6^A-mediated RNA metabolism has not only expanded the discovery of other constituents of the m^6^A-methylation complex, but also the additional RNA substrates, such as lncRNA, that can carry m^6^A modifications. Among these constituents are RBM15 and RBM15B, which have been shown to mediate m^6^A formation in lncRNA *Xist*^116^. Given our identification of RBM15 and RBM15B as enriched VIR-CLASP proteins with robust proviral effects, we investigated whether CedV RNA transcription and replication was dependent on m^6^A methyltransferase activity.

We determined that METTL3-inhibition using small molecule inhibitor STM2457 significantly reduced levels of viral mRNA, cRNA, and new gRNA (Figure 5A). These results were independent of any notable effect on cell viability due to inhibitor treatment (Figure S5A). In addition to methyltransferases, m^6^A-mediated RNA metabolism is also influenced by m^6^A readers. Although there is an expanding list of m^6^A RBPs, we focused on investigating YTH-domain containing family proteins (YTHDF1-3), which have been well characterized for their various functional roles on cellular and viral RNA^25,54,113,117–119^. Thus, we examined whether YTHDF proteins might influence CedV transcription or replication. We did not initially identify YTH family proteins in our VIR-CLASP dataset, which is not too surprising as the labeled incoming genome is negative sense; we would anticipate YTHDF proteins to associate with mRNA transcripts. Nevertheless, if CedV mRNA transcripts contain m^6^A modifications, then it is conceivable that m^6^A readers could influence their fate, given our results with METTL3 inhibition (Figure 5A). We observed that loss of YTHDF 1-3 differentially affected viral RNA. Specifically, depletion of YTHDF1 restricted gRNA and resulting viral production, despite increased expression of cRNA (Figure 5B-C). Loss of YTHDF3 decreased both gRNA and cRNA and reduced nascent virion production (Figure 5B-C). Therefore, YTHDF1 and YTHDF3 are exerting differing effects as required host factors. No significant effects were observed upon YTHDF2 knockdown (Figure 5B-C). Overall, our results demonstrate that CedV transcription and replication is dependent on m^6^A binding proteins and methyltransferases, which suggest that the CedV genome may be m^6^A modified.

**Figure 5.**
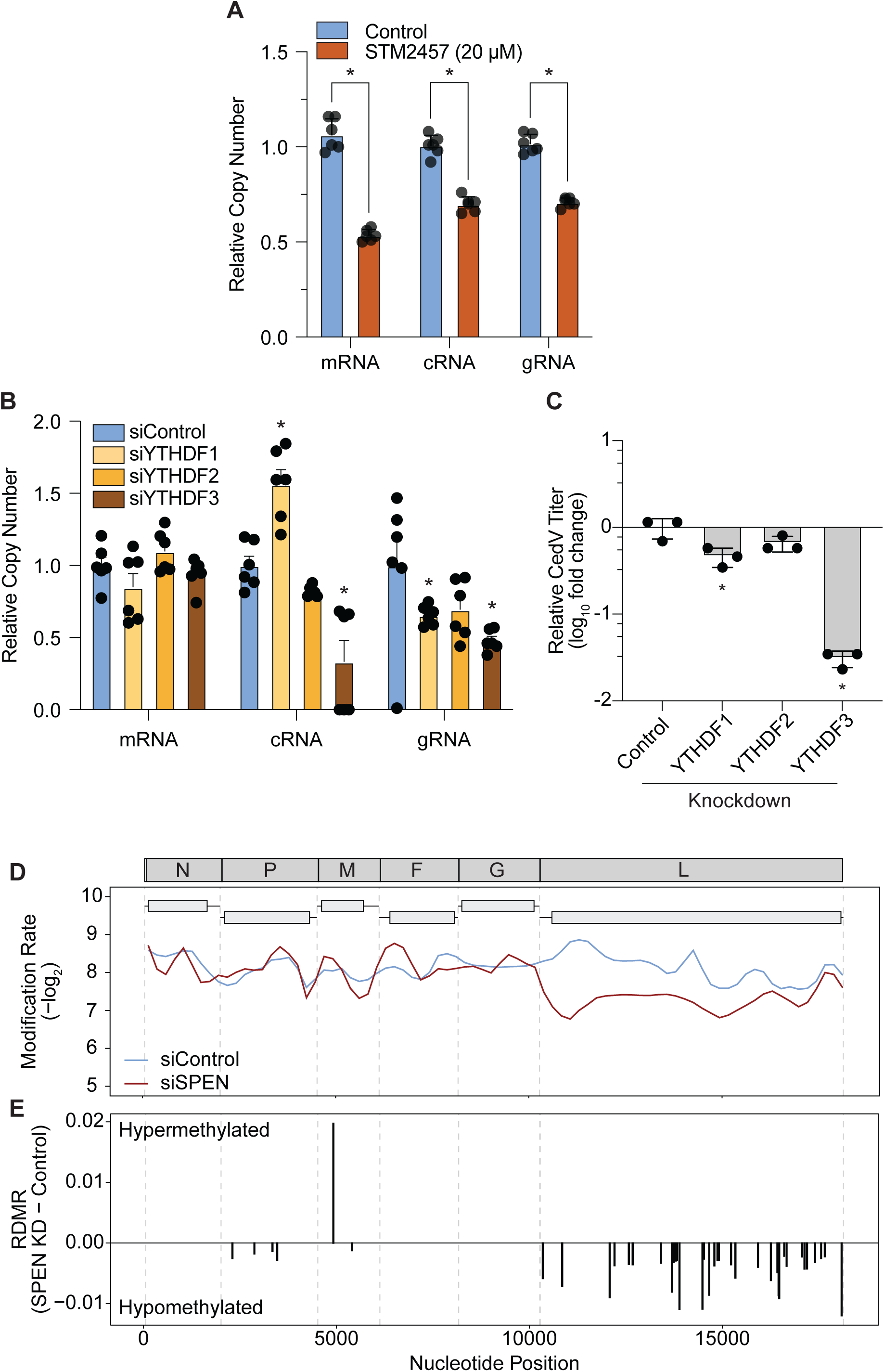
m^6^A modifications in CedV L mRNA are hypomethylated following SPEN depletion. (A) Strand-specific RT-qPCR of HEK293 cells treated with STM2457 (20 µM) and infected with CedV. Cells were treated with STM2457 or DMSO control for 48 h followed by CedV infection (MOI 0.1) for 16 h. n = 6, error bars, mean ± SD. P values were calculated using two-way Anova with Dunnett’s multiple comparisons test. (B) siRNA knockdown of YTHDF-1, -2, -3, or control in HEK293 cells followed by strand-specific RT-qPCR. siRNA transfection was performed for 72 h followed by CedV infection (MOI 0.1) for 16 h. Data is normalized to non-template control. n = 6, error bars, mean ± SEM. P values were calculated using ordinary two-way Anova with Dunnett’s multiple comparisons test. (C) siRNA knockdown of YTHDF-1, -2, -3, or control in HEK293 cells followed by measurement of CedV titer. siRNA transfection was performed for 72 h followed by CedV infection (MOI 0.1) for 16 h. Titer was determined by plaque assay in Vero cells. Data is normalized to non-template control. n = 3, error bars, mean ± SD. P values were calculated using one-way Anova with Dunnett’s multiple comparisons test. (D) Metagene analysis of modification rates in CedV mRNA transcripts from control or SPEN knockdown HEK293 cells. Dark grey schematic represents CedV genome. Alternating light grey boxes represent discrete mRNA transcripts with CDS (light grey rectangles) and UTR regions (black lines). Grey dashed lines represent nucleotide positions of gene start and gene end sites. n = 2 biological replicates. (E) Barplot of relative differential modification rates (RDMRs) in CedV mRNA transcripts. RDMRs were calculated from the difference of modification rates between conditions (SPEN KD – Control). FDR was calculated using a two-tailed, unpooled z-test with Benjamini-Hochberg correction. (+) RDMRs are classified as hypermethylated. (-) RDMRs are classified as hypomethylated. Data represents sites of significant RDMRs (FDR < 0.05). n = 2 biological replicates

### m^6^A modifications in CedV L mRNA are hypomethylated following SPEN depletion

Previous studies have shown that m^6^A formation in *Xist* is mediated by RBM15 and RBM15B^120,121^. Specifically, RBM15/15B were found to promote *Xist* methylation through the recruitment of WTAP-METTL3 complexes and that knockdown of both RBM15 and RBM15B reduced m^6^A levels^116^. While RBM15/15B have been characterized as additional components of the m^6^A methyltransferase complex^120,122^, the involvement of SPEN in m^6^A modifications has yet to be investigated. Intriguingly, METTL3 has been captured in the protein interactome of the SPEN SPOC-domain^123^. Our findings, together with the established m^6^A regulatory roles of RBM15 and RBM15B, suggested a model in which SPEN – unlinked to m^6^A modification – similarly directs methylation of CedV mRNAs.

We performed DRS to profile the m^6^A modification landscape in CedV mRNA following SPEN depletion. The CedV genome contains six gene segments that synthesize RdRp-mediated capped and polyadenylated mRNA encoding for the six structural proteins^11^. To capture these mature viral mRNA transcripts, we first collected total RNA from CedV-infected siControl and siSPEN HEK293 at 16 hpi and prepared sequencing libraries from the poly(A)+ RNA fraction. Following raw data analysis using Dorado basecalling model^124^ and Minimap2 alignment^125^, we observed ~10% and ~5% of mapped reads aligned to the viral reference sequence in Control and siSPEN samples, respectively (Figure S5C). These results are not only consistent across biological replicates but are also in agreement with the MOI (MOI 0.1) used to infect the cells and the reduction in viral mRNA we consistently observe upon SPEN knockdown. To computationally identify m^6^A sites for downstream analyses, we used Modkit for single site analysis of modified bases with a canonical A base threshold ≥ 0.9 and base modification threshold ≥ 0.7. We also filtered for sites matched across biological replicates and sample conditions and regions with coverage greater than 50 identified unmodified or modified adenosines. We calculated the proportion of m^6^A-modified adenosines at a single site and define these values as the modification rate for a given nucleotide position. This computational approach enables statistically robust m^6^A detection and quantification of differentially modified sites between control and SPEN knockdown samples. We also determined that modification rates between biological replicates were highly correlated (Figure S5D-E).

Overall, we identified 441 putative m^6^A sites with varying modification rates in both samples (Table S4). Metagene analysis of m^6^A sites revealed a broad distribution of modifications detected within the CedV genome from control samples (Figure 5D). Further, we broadly observe elevated modification rates in the CDS regions of corresponding mRNA transcripts. Of note, we found that m^6^A modifications roughly localized near UTR/CDS boundaries although further investigation is warranted. Our analysis also resolved differentially modified regions that were either hyper- or hypomethylated following SPEN depletion (Figure 5D). Our results of CedV m^6^A profiles in control conditions are supported by recent work that used DRS to determine the m^6^A state of two strains of NiV^114^.

We next mapped sites differentially modified upon SPEN depletion by calculating the relative differential modification rate (RDMR), which we defined as the difference in modification rates (MR) between SPEN-depleted and control conditions (MR_SPEN KD_ – MR_Control_)^126^ at a single nucleotide position. Each RDMR was classified as hypermethylated (+ RDMR) or hypomethylated (-RDMR), and statistical significance was assessed by FDR. Applying an FDR threshold of < 0.05, we identified 40 sites with significant differential m^6^A modification (Table S4). Remarkably, the overwhelming majority (~98%, 39/40 sites) were hypomethylated, supporting the hypothesis that SPEN depletion reduces m^6^A deposition. Even more striking was the location of these modifications. ~87% (34/39 sites) of these hypomethylated sites were clustered within the L mRNA, which encodes the viral RdRp essential for viral transcription and replication (Figure 5E and Table S4). Overall, the predominant hypomethylation of significant differential m^6^A sites – clustered almost exclusively within the L mRNA – reveals that SPEN exerts highly selective control over m^6^A deposition. By fine-tuning post-transcriptional regulation of the RdRp, the core viral component of the replication-transcriptional complex, SPEN orchestrates the earliest and most consequential events of viral genome replication, providing a key mechanistic insight into a poorly characterized layer of host-mediated control over henipavirus infection.

## DISCUSSION

Interrogation of primary interactions between vRNA and host RBP provides crucial insight into vRNA regulation during early infection. In this study, we provide molecular evidence for pioneer determinants of henipavirus pathogenesis, which is a rapidly emerging group of single stranded RNA viruses including BSL-4 members, HeV and NiV. Using CedV as a tractable platform to model henipavirus infection, we applied the photocrosslinking technique, VIR-CLASP with genetic perturbation experiments and direct RNA sequencing to define the pre-replicated CedV RNA interactome and reveal functional roles of vRNA interactants in shaping viral infection. We identified 146 cellular proteins directly bound to the incoming CedV genome and further characterized SPEN family proteins - SPEN, RBM15, and RBM15B - as crucial proviral host factors. Upon depletion of SPEN, we observed a striking reduction of m^6^A modification, ~98%, with remarkable selectivity; ~87% were localized in the L mRNA transcript. Our results highlight a previously unknown role for SPEN as a regulator of m^6^A deposition, and an unappreciated role for at least three SPEN family proteins as immediate-early determinants in regulating early viral infection.

SPEN, originally characterized as a transcriptional regulator and *Xist* lncRNA binding protein, emerged as a direct interactant of vRNA and robust proviral factor. Although SPEN has appeared in previous viral interactome datasets^26,104^, its biological significance during viral infection has remained unresolved. Our work establishes a direct and functional role for SPEN in promoting infectious virion production, viral transcription, and replication. These findings elevate SPEN from a recurrent interactor to a *bona fide* proviral determinant, revealing mechanistic links between SPEN activity and vRNA regulation. In addition to uncovering SPEN, we captured RBM15 and RBM15B as enriched vRNA interactants, also consistent with prior identifications in vRNA interactomes^26,33,37,104^. Like SPEN, these proteins had previously uncharacterized roles in viral infection. Our data suggest that RBM15 and RBM15B also function as proviral factors. It should be noted that our study also identified PHF3, another SPOC-domain protein. PHF3 bridges Pol II elongation machinery with chromatin and RNA processing factors^127,128^, suggesting that SPOC-domain proteins may broadly scaffold transcriptional and post-transcriptional regulatory complexes on vRNA. Our data further hint at what appears to be an epistatic relationship between SPEN and RBM15/15B, raising the possibility that these proteins partially compensate for, or even compete during vRNP assembly and transcription. Alternatively, RBM15/15B may direct m^6^A modifications in partially non-overlapping mRNA sites relative to SPEN, resulting in alterations in the fates of the other CedV mRNAs – all of which were found to contain m^6^A modification. The precise dynamics of this interplay during viral infection merits further investigation.

In the context of cellular RNA, m^6^A sites within the CDS typically affect transcript stability and abundance relative to other regions by triggering translation-dependent mRNA decay^129^. It remains unclear how the location of m^6^A sites in viral mRNA influences stability and translation. The dramatic decline of m^6^A modification rates in the CedV L mRNA UTR-CDS boundaries suggests a potentially location-dependent regulatory mechanism facilitated by SPEN. While we use CedV as a tractable model for henipavirus infection, it would be interesting to evaluate how SPEN influences the m^6^A landscape in HeV and NiV and whether loss of SPEN selectively remodels certain mRNA transcripts, particularly V and W protein, which are accessory proteins and immune antagonists that are not encoded by CedV. Further, we speculate that selective hypomethylation of the L mRNA, and thus potentially limited protein production of the RdRp, may be consequential to the synthesis of other CedV mRNA transcripts, which we observe when we examine the expression of other CedV mRNA transcripts following SPEN depletion. Given the high virulence of HeV and NiV^4^ and recent identification of m^6^A modifications in NiV RNA^114^, it is attractive to speculate that the pathogenicity of these BSL-4 agents can be linked to distinct m^6^A landscapes that are remodeled by pioneer host factors.

Taken together, our findings favor a model in which SPEN, together with other SPEN family proteins, are recruited and directly bound to the incoming vRNA genome. Upon recruitment, they are able to help nucleate methyltransferase machinery to modulate m^6^A modifications on subsequent strands of viral RNA. Given our current data, it is unclear whether the functional activity of SPEN is dependent on or associated with the viral RdRp. Previous structural studies have shown that SPEN and RBM15 SPOC domain serve as reader domains mediating phosphoserine binding on the CTD of RNA Pol II^130^. It is unclear whether these features are conserved on viral RdRp and whether these binding properties are crucial to SPEN activity on vRNA. If the viral RdRp harbors similar binding modules, it is conceivable that SPEN activity may be mediated by viral RdRp association. Nonetheless, if SPEN were to physically associate with viral RdRp, it does so in a manner that appears more selective as the loss of SPEN had a profound reduction of m^6^A principally within the L mRNA. It is possible that SPEN can interact with the incoming RdRp complex, pre-packaged within the incoming virion, in a distinct fashion than it does with new RdRp complexes that are synthesized during infection.

We define the pre-replicated CedV RNA interactome to consist of 146 distinct cellular proteins. In this report, we evaluated a subset of candidates, which we categorize as either antiviral or proviral factors in CedV infection. While we characterize the functional role of SPEN family proteins, we anticipate that our interactome is comprised of other crucial regulators of vRNA metabolism and viral infection. Among our identified antiviral host factors, we determined that loss of WDR33 increased nascent virion production (Figure 2D). As a constituent of the pre-mRNA 3’ processing complex^70^, we speculate a potentially uncovered function in which WDR33 may be engaged to the incoming CedV genome to influence pre-mRNA maturation and destabilize viral transcription through regulation of 3’ end cleavage and poly(A) tail synthesis. Additionally, we uncovered several proviral factors, such as splicing factor and nuclear speckle driver, SRRM2^78,79^ (Figure 2D). Other RNA viruses, such as IAV, have been characterized to co-opt splicing factors to produce multiple viral transcripts from one gene segment during viral replication^131,132^, raising the intriguing possibility that CedV association with SRRM2 leads to alternative splicing events or other post-transcriptional processes that might require localization to nuclear speckles. While the role of splicing in henipavirus infection has yet to be characterized, we speculate that viral transcription may require cellular splicing factors, such as SRRM2, to facilitate the synthesis of viral transcripts, particularly for mRNA encoding accessory proteins, V/W/C, that are alternative products of the P gene^11^. Since these proteins are produced through an alternate transcriptional mechanism of mRNA editing or use of an alternate open reading frame^11^, it is attractive to speculate that host splicing factors, such as SRRM2, are hijacked during this critical viral process.

Overall, this study reveals the vRNP-RBP dynamics and post-transcriptional circuitry that orchestrate henipavirus infection, providing a mechanistic foundation that elevates our understanding of an otherwise inaccessible genus of RNA viruses. Our work spotlights SPEN, RBM15, and RBM15B as attractive host targets, resistant to viral evolution, owing to their dependency by the vRNA to install cellular post-transcriptional marks and thus opens new avenues for the rational design of antiviral therapeutics.

### Limitations of the study

While we show that SPEN differentially affects m^6^A sites in viral mRNA, we cannot rule out the possibility that the role of SPEN in regulating viral RNA may include other post-transcriptional mechanisms beyond modulating m^6^A modifications. Previous studies capturing the protein interactome of the SPEN SPOC domain have uncovered a broad array of co- and post-transcriptional regulators that are functionally associated with RNA Pol II activity and have posited SPEN as a molecular integrator for transcriptional machinery on cellular RNA^123,128,130^. Thus, given the widespread interactions of proteins partners with the SPOC domain^123,127^, it is conceivable that SPEN may additionally scaffold other co- or post-transcriptional regulatory effectors in vRNP assembly and may not be exclusive to bridging m^6^A machinery.

## RESOURCE AVAILABILITY

### Lead contact

Manuel Ascano

### Materials availability

Further information and requests for resources and reagents should be directed to and will be fulfilled by the lead contact, Manuel Ascano.

### Data and code availability

Mass spectrometry raw data have been deposited to the ProteomeXchange Consortium (<u>Deutsch et al.,</u> <u>2017</u>) via the PRIDE (<u>Perez-Riverol et al., 2019</u>) partner repository with the dataset identifier PRIDE: *in progress*. Nanopore direct RNA sequencing raw data have been deposited to National Center for Biotechnology Information via Sequence Read Archive (SRA); BioProject ID: *in progress.* All code used for proteomic analysis and figure generation is accessible at https://github.com/Ascano-Lab.

## ACKNOWLEDGEMENTS

We would like to thank Drs. C. C. Broder and M. Amaya for providing CedV and CedV-GFP; the Mass Spectrometry Research Center Proteomics Core Laboratory (Vanderbilt University), in particular Dr. K L Rose and Dr. M. Leser; Dr. C. P. Plamen and Vanderbilt Institute of Chemical Biology Chemical Synthesis Core for 4SU synthesis; Dr. L. Plate for discussion in the MS analysis; Vanderbilt Medical Center Flow Cytometry Core, in particular D. K. Flaherty, B. K. Matlock, O. X. Murfield, and E. McLaughlin for flow cytometry analysis; Drs. J. Karijolich and Dr. X. Ye for technical discussion and assistance in direct RNA sequencing. Finally, we would like to thank members of the Ascano laboratory for their support, collegiality, and critical review of the manuscript. Flow Cytometry experiments were performed in the VMC Flow Cytometry Shared Resource. The VMC Flow Cytometry Shared Resource is supported by the Vanderbilt Ingram Cancer Center (P30 CA68485) and the Vanderbilt Digestive Disease Research Center (DK058404). This work was supported by the NIH (5R35GM119569 to M.A.) and Cellular, Biochemical, and Molecular Sciences Training Program (training grant 3T32GM137793 to A.A.)

## AUTHOR CONTRIBUTIONS

Conceptualization & methodology, AA, SC, JT, JMC, and MA; Software and Data Curation, AA and SC; Analysis, AA and MA; Investigation, AA, AL, and SH; writing—original draft, AA; writing—review & editing, AA and MA; funding acquisition, MA; supervision, MA.

## SUPPLEMENTAL INFORMATION

**Document S1. Figures S1–S5.**

**Table S1. Proteomics Analysis from VIR-CLASP with CedV, Related to Figure 1.**

**Table S2. List of putative and previously identified RBPs based on published vRNA interactomes, Related to Supplementary Figure 1.**

**Table S3. Functional enrichment analyses of the CedV RNA interactome, Related to Figure 2.**

**Table S4. Nanopore DRS analysis of m^6^A sites and relative differential modification rates in CedV mRNA transcripts, Related to Figure 5.**

**Table S5. Primers and siRNAs used in this study, related to the STAR Methods.**

## METHODS

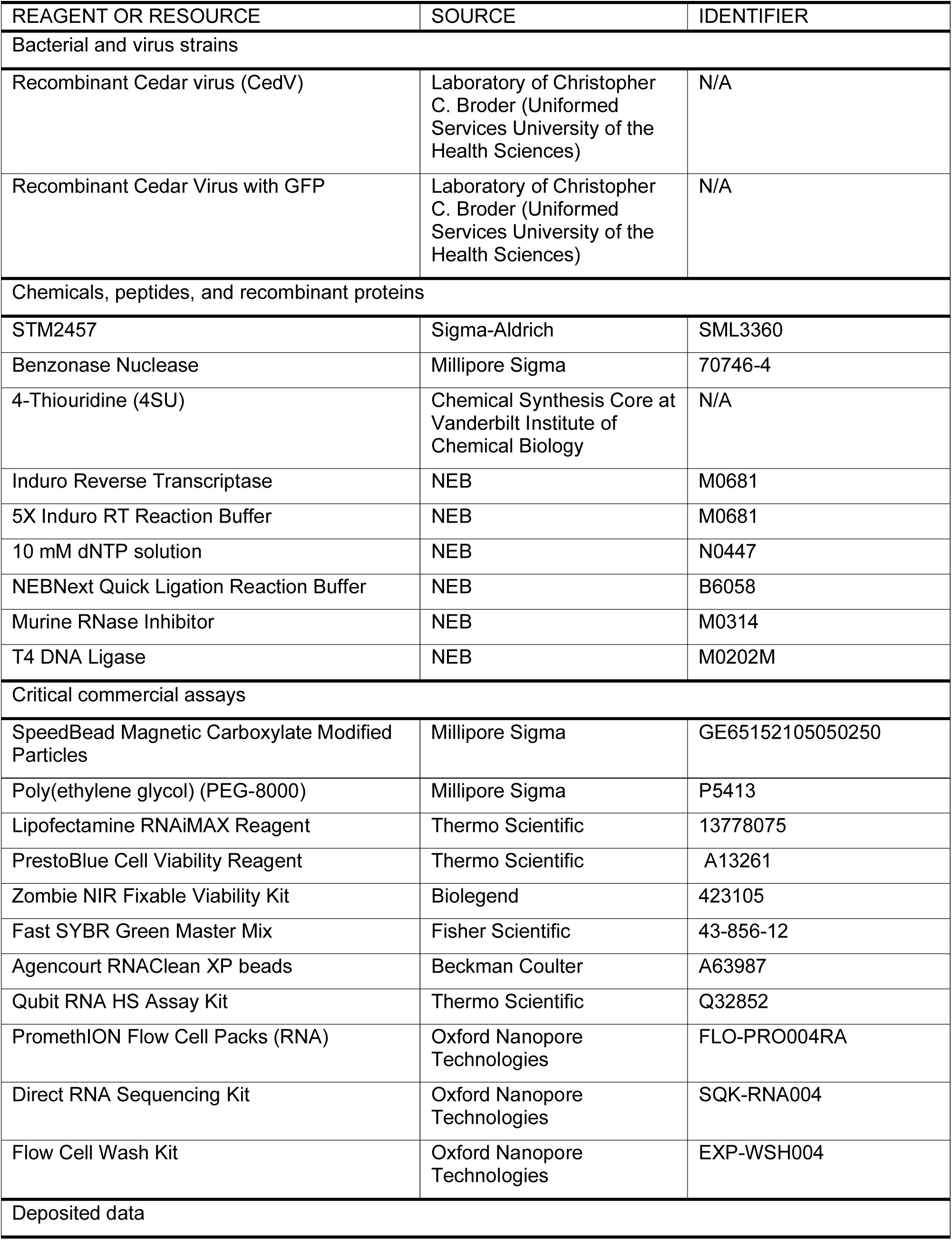
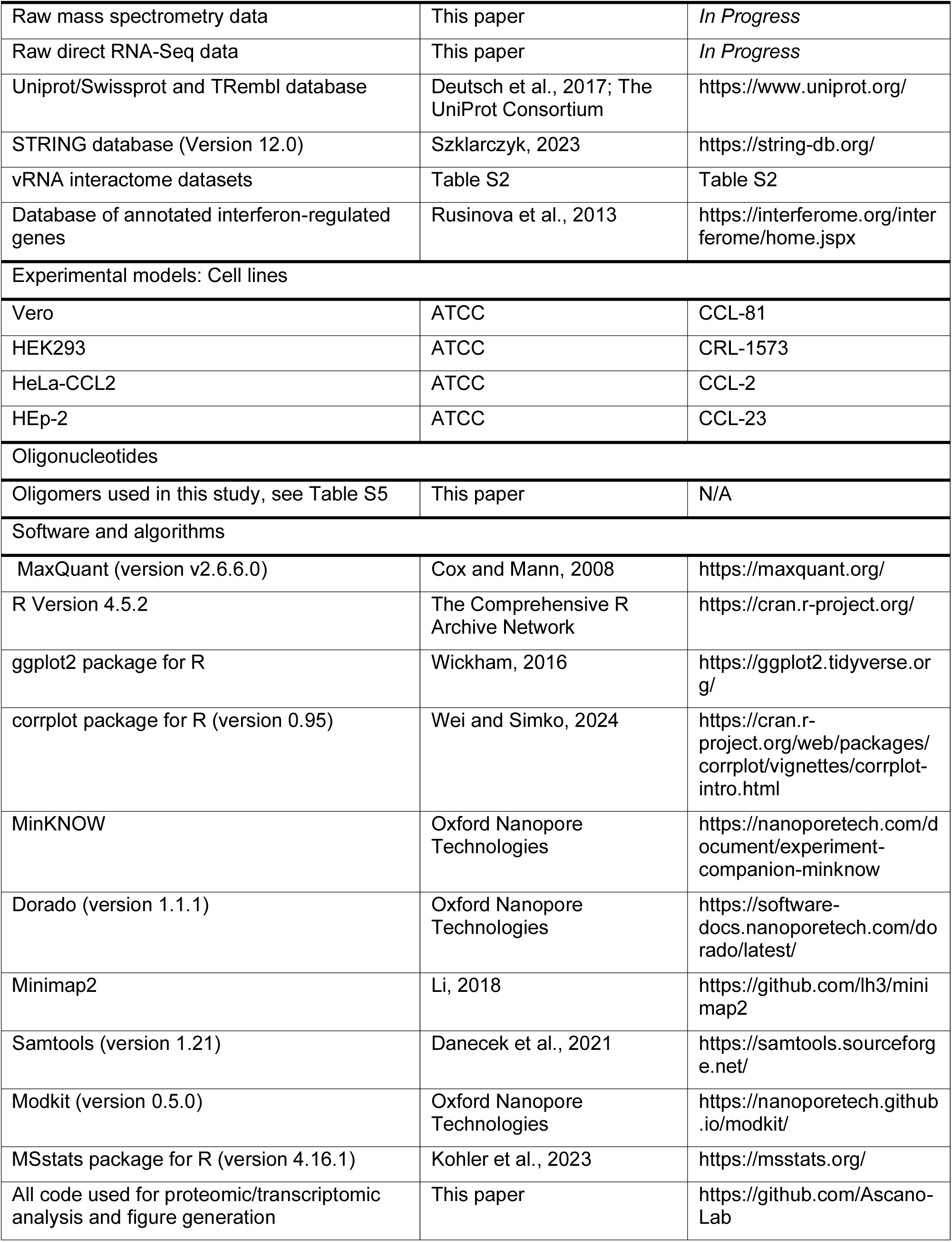
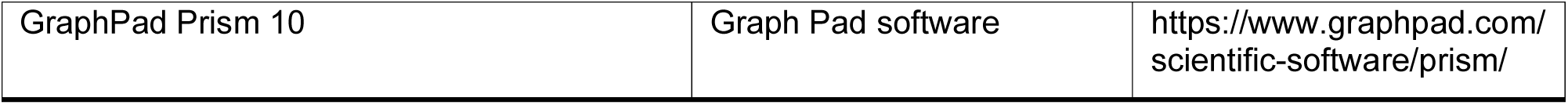

### EXPERIMENTAL MODEL

#### Cell lines and culture

Cell lines were obtained from ATCC. Vero cells (female) and HEK293 cells (female) were maintained in DMEM (Gibco) supplemented with 10% fetal bovine serum (FBS from Peak Serum) and 2mM L-Glutamine (Gibco) and 100 U/mL penicillin/streptomycin (Gibco).

#### Viruses

Recombinant Cedar virus was provided by Christopher C. Broder (Uniformed Services University). CedV was propagated in Vero cells with or without 1 mM 4-thiouridine (4SU). Virus stocks were purified by two rounds of ultracentrifugation of clarified supernatants through a 20% sucrose cushion in TNE buffer (20mM Tris-HCl, 150mM NaCl, 1mM EDTA, pH 7.4) at 100,000 x g for 2 h in a Beckman SW32Ti rotor. Virus pellets were washed with TNE buffer to remove remaining free 4SU, re-suspended in 20% sucrose in TNE buffer, and stored at −80°C. Virus titers were determined by plaque assay using Vero cells.

### METHOD DETAILS

#### VIR-CLASP

##### Viral Infection and UV365nm crosslinking

For VIR-CLASP, cells were infected with MOI 1000 of 4SU-labeled virus for 1 hr at 37°C and uninfected virus was washed away with PBS. The infected cells were incubated for an additional 1 hr at 37°C prior to 365 nm ultraviolet irradiation. Prior to UV365nm irradiation, the growth medium was removed, and cells were washed with PBS. Cells were irradiated on ice with 365 nm UV light (0.6 J/cm2 x 3 times) in a Stratalinker 2400 (Stratagene). Cells were scraped off in 2.5 ml PBS per plate.

##### CLASP

Cells were lysed in denaturation buffer (50 mM Tris–HCl, pH 6.8, 10% glycerol, 2.5% SDS, 0.66% NP-40), incubated for 10 min at 95 °C and subsequently slowly cooled to 25°C. Crosslinked RNA-protein complexes were purified by Solid-Phase Reversible Immobilization (SPRI) (Hawkins et al., 1994) beads (GE Healthcare, cat# 65152105050250) under denaturing SPRI buffer (30 mM Tris–HCl, pH 6.8, 6% glycerol, 1.5% SDS, 0.4% NP-40, 1 M NaCl, 8% PEG-8000). To each sample, 0.66× (e.g. 660 μl of beads for 1ml of sample) of SPRI beads (1mg/ml SPRI beads in 10 mM Tris-HCl, pH 8.0, 1 M NaCl, 18% PEG-8000, 1 mM EDTA and 0.055% Tween 20) were added, and samples were incubated at room temperature for 10 minutes. The SPRI beads and complexes were washed 5 times with denaturing SPRI buffer. The crosslinked RNA-protein complexes were eluted for 5 min at 37 °C in denaturation buffer (lysis buffer). To reduce non-specific binding on the beads, SPRI purification was repeated. To digest RNA from crosslinked RNA-protein complexes, an equal volume of 4x Benzonase buffer (80 mM Tris-HCl, pH 7.5, 600 mM NaCl, 20 mM MgCl2, 4 mM DTT, 40% Glycerol) and 2x volume of water were added to eluted samples, followed by the addition of Benzonase (EMD Millipore, cat# 70746-4) to a final concentration of 50 U/ml, and incubation for 2 hr at 37°C. Proteins were precipitated by methanol and chloroform and then re-suspended in 2x NuPAGE LDS Sample Buffer (Thermo Fisher Scientific, cat# NP0007) with 50 mM DTT or 100mM TEAB for silver staining or mass spectrometry analysis, respectively.

#### Plaque Assay

Vero cells were seeded on 12-well plates (2 x 10^5^ cells/well) one day prior to infection. Next day, infectious supernatant was serially diluted using 10-fold dilutions in DMEM (Gibco) supplemented with 10% FBS and used to infect Vero cells for 1 h at 37°C and 5% CO2, rocking the plate every 15 minutes. After infection, cells were overlaid with carboxymethylcellulose overlay media (1:1 mix of DMEM supplemented with 5% FBS and 2% Carboxymethylcellulose (CMC) (MP Biomedicals, cat# 150560) was incubated at 37°C and 5% CO2 for five days. Cells were then fixed with 10% formaldehyde and stained with 0.5% crystal violet. The resulting plaques were counted.

#### Mass Spectrometric Analysis

Protein samples were brought to 5% SDS, prepared by S-Trap (ProtiFi) digestion, and analyzed by LC-coupled tandem mass spectrometry (LC-MS/MS) similar to methods previously described^133^. Proteins were initially reduced with 10mM TCEP, alkylated with 20mM iodoacetamide, and aqueous phosphoric acid was added at a final concentration of 2.5% followed by S-trap binding buffer (90% methanol in 100 mM TEAB) at 6 times the sample volume. Samples were loaded on S-Trap micro columns, washed with S-trap binding buffer, and were digested with1ug trypsin gold (Promega) in 50 mM TEAB, pH 8.0, for 1 hr at 47°C. Peptides were eluted by serial addition of 40 µL each of 50 mM TEAB, 0.2% formic acid, and 35 µL of 0.2% formic acid in 50% acetonitrile. Eluted peptides were dried and resuspended in aqueous 0.2% formic acid. For LC-MS/MS, peptides were loaded onto a C18 reverse phase analytical column using a Dionex Ultimate 3000 nanoLC and autosampler and were gradient eluted using a 90 min gradient. The gradient consisted of the following: 1-77 min, 2–38% B; 77-82 min, 38–95% B; 82–83 min, 95% B; 83-84 min, 95–2% B; 84-90 min (column re-equilibration), 2% B. Peptides were analyzed using a data-dependent method on an Orbitrap Exploris 240 mass spectrometer (Thermo Scientific), equipped with a nanoelectrospray ionization source. The instrument method consisted of MS1 using an MS AGC target value of 3 × 106, followed by 20 MS/MS scans of the most abundant ions detected in the preceding MS scan. The intensity threshold for triggering data-dependent scans was set to 1 × 104, the MS2 AGC target was set to 1 × 105, dynamic exclusion was enabled (15sec), and HCD collision energy was 30 nce.

#### Bioinformatics analysis of Mass Spectrometry data

Raw data files from the LC-MS/MS instrument were processed and searched using MaxQuant (v2.5.2.0)^134^ to generate peak lists and identify peptide-spectrum matches. Searches were performed using a Uniprot/Swissprot database for *Homo sapiens* (Proteome ID: UP000005640), *Chlorocebus sabaeus* (Proteome ID: UP000029965), and *Cedar virus* (Proteome ID: UP000116133) including both reviewed (Swiss-Prot) and unreviewed (TrEMBL) proteins (downloaded on 26 May 2024), with added sequences for Benzonase nuclease (Uniprot Accession: P13717). The search parameters for Andromeda included full tryptic specificity, two missed cleavages allowed, carbamidomethyl (C) fixed modification, and acetylation (N terminal) variable modification. Match between runs was selected, and LFQ normalization with iBAQ identification algorithm. All other settings used were default, resulting in a protein FDR of < 0.01 for each dataset. To define pre-replicated CedV interacting proteins, we used the MSstats R package (v4.16.1).^135^ Parameters for MSstats included the following: Potential contaminants were retained, proteins identified by only one peptide were removed, and protein identification was performed using at least unique peptides. Quantile normalization was performed to remove systematic bias between MS runs, model-based imputation was performed for censored missing values using accelerated failure time model, and remaining parameters were set to default. For statistical quantification of differentially abundant proteins, +4SU CLASP sample intensities were compared to the corresponding +4SU Input sample intensities. Adjusted P values were computed using Student’s t-test with Benjamini-Hochberg correction. Human UniProt/Swiss-Prot proteins with an adjusted p-value ≤ 0.05 and a log_2_ fold-change > 1 were defined as enriched VIR-CLASP proteins (CedV RNA interactome). Correlations between sample replicates were determined using Pearson correlation.

#### Functional enrichment analyses

Functional enrichment clusters of Gene Ontology terms in protein-protein association network were retrieved and clustered STRING v12.0 database. Enriched Reactome pathways and InterPro classifications were queried using STRING v12.0 database and applying default settings. Statistical background for enrichment analysis was performed using the whole genome.

#### siRNA knockdown

siRNAs used in this study are listed in Table S5. HEK293 cells were seeded in 24-well plates at 1.5 x 10^5^ cells/well and incubated 24 h prior to transfection. siRNAs were transfected at a final concentration of 20nM using Lipofectamine® RNAiMAX (Invitrogen) according to the manufacturer’s instructions. Cells were then incubated at 37°C and 5% CO2 for 72 h prior to viral infection with the indicated MOI.

#### STM2457 treatment

Small molecule inhibitor, STM2457 (Sigma-Aldrich, cat no. SML3360) was reconstituted in DMSO as a 10 mM stock solution and subsequently diluted in infection media prior to cell treatment. STM2457 stocks were stored at −80°C. For inhibitor treatments, cells were incubated in 20 µM STM2457 for 48 h, then infected with CedV (MOI 0.1). Cell lysates were collected 16 hpi in Trizol for RNA extraction.

#### Cell viability assay

HEK293 cells were seeded in 96-well plates at 1.5 x 105 and treated with STM2457 inhibitor at varying concentrations. Stock solutions were diluted in infection media prior to cell treatment. DMSO was used as a vehicle control. After 48 h of treatment, cell viability was assessed using PrestoBlue reagent (Thermo Scientific, cat no. A13261) according to the manufacturer’s instructions.

#### RNA extraction and RT-qPCR analysis

RNA was collected and extracted from infected or uninfected cells using Trizol (Ambion). The total RNA concentration was determined using a NanoDrop 8000 (ThermoFisher). 1 µg total RNA for each sample were reverse transcribed using SuperScript III (ThermoFisher) with oligo(dT) primers or CedV strand-specific primers. Real-time PCR reactions were performed using FastSYBR Green Plus Master Mix (Applied Biosystems) using a QuantStudio™ 3 Real-Time PCR instrument (Applied Biosystems). Oligonucleotides used in this study are listed in Table S5. For relative quantification, target Ct values were normalized to TUBA1A Ct values and used to calculate ΔCt. Relative mRNA expression of target genes was then calculated using the ΔΔCt method. For absolute quantification of viral copy number, a standard curve was generated using a plasmid containing a cDNA copy of a region of the CedV P gene. Viral copy number was calculated based on target Ct values using the standard curve and relative viral copy number was determined based on control (NTC) values.

#### Flow Cytometry

For GFP expression analysis, 8 x 10^5^ HEK293 cells were seeded on 6-well plates and transfected with siRNA using method outlined above. Cells were then infected with or without CedV-GFP at MOI 0.1 for 1 hour at 37°C, followed by aspirating the virus inoculum, washing with 1X PBS, adding DMEM containing 10% FBS and 1% L-glutamine, and incubating at 37°C for the indicated timepoint. At the indicated hpi, the cells were washed with 1X PBS, collected with Accutase (Sigma-Aldrich, cat no. A6964), transferred to tubes, and washed with 1X PBS. Cells were stained live with Zombie NIR viability dye (Biolegend). After staining, cells were washed with FACS buffer (1X PBS containing 1% BSA and 5mM EDTA) and fixed in 4% PFA for 20 mins at 4°C. PFA was aspirated, and cells were resuspended in FACS buffer for flow cytometry analysis. For every sample, at least 30,000 live events were analyzed on a BD 3-Laser Fortessa. Quantification of GFP expression was gated based on mock-infected condition for each experiment. Data were analyzed with FlowJo and plotted with Floreada.io.

#### Nanopore Direct RNA sequencing and bioinformatic analysis

Direct RNA-sequencing libraries were generated from 1 µg total RNA and RNA integrity was evaluated using RNA ScreenTape analysis (Agilent, cat no. 5067-5576; 4150 TapeStation system) prior to library preparation. Sequencing libraries were prepared using Oxford Nanopore Technologies (ONT) Direct RNA sequencing kit (SQK-RNA004) and DRS protocol (DRS_9195_v4_revI_30Jul2025). Standard reverse transcription adapters and T4 DNA Ligase was used to capture poly(A)+ fraction, and cDNA was synthesized using Induro reverse transcriptase to improve RNA sequencing output. Libraries were sequenced on PromethION S2 with flow cells (FLO-PRO004RA). Following sequencing, basecalling was performed on POD5 files using Oxford Dorado simplex basecalling model with modified bases for m^6^A in a DRACH context (sup@v5.2.0,m6A_DRACH@v1) with a mean Q-score 7. Only reads present in the “pass” folder were used in subsequent analyses. Basecalled read data were aligned to the CedV isolate CG1a reference genome (NCBI RefSeq: GCF_000924595.1) and *Homo sapiens* Hg38 genome using Minimap2 aligner with default presets^125^. Aligned BAM files were sorted and indexed using SAMtools v1.19.2.^136^ Following alignment, Modkit (https://nanoporetech.github.io/modkit/) was used for subsequent modified base analysis. Modkit pileup was used to generate bedMethyl tables containing counts of modified and unmodified bases from every sequencing read over each aligned genomic position with a canonical ‘A’ threshold > 0.9 and base modification threshold > 0.7. Only primary alignments were used for unmodified and modified base counts. Modkit dmr pair command was used to calculate max coverage from read data and perform differential methylation scoring at single nucleotide positions across conditions. Sites were further filtered for those matched across biological replicates and sample conditions and with valid coverage > 50 total unmodified or modified adenosines. These filtered sites are used for downstream analysis of differential modifications. The modification rate (MR) for a given nucleotide position for each condition is quantified as the proportion of m^6^A-modified adenosines relative to the total number of adenosines (m^6^A/A) at a single site. Using the modification rates for each sample, we calculated a relative differential modification rate (RDMR), which we define as the difference between modification rates of each condition (MR_SPEN KD_ – MR_Control_). The directionality of the RDMR is classified as either a hypermethylated m^6^A site (+ RDMR) or hypomethylated m^6^A site (-RDMR). To determine statistically significant sites with differential modification rates in a pairwise comparison, we performed a two-tailed, unpooled z-tests on the modification rate difference of control and SPEN knockdown conditions for each nucleotide position, using the average of valid read coverages and modification rates across biological replicates for each condition^126^. Resulting z-scores were used to rank differentially modified positions and compute corresponding P-values, which were adjusted using Benjamini-Hochberg correction. Statistically significant differential modification rates are defined as those with FDR < 0.05. Data visualization of modification rates and differential modification rates were made with R package ggplot2^137^ using CedV genomic boundaries and CDS regions (GenBank: JQ001776.1) to define localization. All code for the Nanopore DRS data processing and R analysis can be found at https://github.com/Ascano-Lab.

### QUANTIFICATION AND STATISTICAL ANALYSIS

Quantification and statistical analyses were performed using GraphPad PRISM 10 software or R programming. Statistical tests, error bars, and numbers of biological replicates of assays (n) are provided and defined within the corresponding figures or figure legends.

