## Supplemental Figures and Legends for "Early Host-Virus RNA Interactions Reveal SPEN-Driven m^6^A Regulation as a Major Determinant of Henipavirus Infection"

### SUPPLEMENTARY FIGURE TITLES AND LEGENDS

**Supplementary Figure 1. Capturing CedV RNA-protein interactions during pioneer events**

(A) RT-qPCR of ephrin-B1 and -B2 expression in uninfected HEK293, HeLa, HEp-2, and Vero cells. Data is normalized to ephrin-B1 or -B2 expression levels in HeLa cells. n = 3, error bars, mean ± SD.

(B) Heatmap and hierarchical clustering of differentially abundant protein intensities identified in Input and CLASP samples. Average protein intensities of replicates are displayed. Scale bar represents Z-score computed for each $\log_2$ protein intensity.

(C) Left: Heatmap and hierarchical clustering of differentially abundant proteins identified between Input replicates. Scale bar represents Z-score computed for each $\log_2$ protein intensity. n = 4. Right: Heatmap and hierarchical clustering of differentially abundant proteins identified between CLASP replicates. Scale bar represents Z-score computed for each $\log_2$ protein intensity. n = 4.

(D) Proportion of CedV RNA interactome that are previously reported RBPs or putative RBPs.

**Supplementary Figure 2. Functional enrichment analysis of the pre-replicated CedV interactome**

(A – B) Dot plot of Reactome pathway enrichment analysis for cellular- (A) and virus-related processes (B). Circles are scaled to gene count in each term. Enrichment is determined from all human proteins. Adjusted P values were calculated using Fisher's exact test with Benjamini-Hochberg correction.

(C) Protein-protein association network of CedV RNA interactome retrieved from STRING v12.0[89]. Colored circles represent closely related GO terms. Colored annotations represent 78% of interactome candidates. Connections represent associations and interactions between proteins.

**Supplementary Figure 3. SPEN is a robust proviral factor necessary for viral infection**

(A) siRNA knockdown of CedV RNA interactome proteins in HEK293 cells followed by RT-qPCR to measure mRNA expression and evaluate knockdown efficiency. Data is normalized to TUBA1A using $\Delta\Delta C_t$ method and is relative to non-template control. n = 3, error bars, mean ± SD. P values were calculated using one-way Anova with Dunnett's multiple comparisons test.

(B) Schematic of strand-specific RT-qPCR primer design to measure gRNA, mRNA, and cRNA. Red arrows represent reverse transcription primers. Dark grey arrows represent PCR primers.

(C – D) Standard curve for absolute quantification of (-) strand gRNA (C) and (+) strand mRNA/cRNA (D) copy number for strand specific qPCR.

**Supplementary Figure 4. Additional SPEN family proteins, RBM15 and RBM15B promote viral infection**

(A) siRNA knockdown of CedV RNA interactome proteins in HEK293 cells followed by RT-qPCR to measure mRNA expression and evaluate knockdown efficiency. Data is normalized to TUBA1A using $\Delta\Delta C_t$ method and is relative to non-template control. n = 3, error bars, mean ± SD. P values were calculated using one-way Anova with Dunnett's multiple comparisons test.

(B) siRNA knockdown combinations of RBM15, RBM15B, SPEN, or control HEK293 followed by infection with CedV-GFP (MOI 0.1) for 12 h. Cells were analyzed by flow cytometry to determine mean fluorescence intensities of GFP. n = 2, error bars, mean ± SD. P values were calculated using ordinary one-way Anova with Dunnett's multiple comparisons test.

(C) siRNA knockdown of RBM15, RBM15B, SPEN, and control in HEK293 cells followed by RT-qPCR of ephrin-B1 and -B2 in uninfected HEK293 cells. Data is normalized to TUBA1A using $\Delta\Delta C_t$ method and is relative to non-template control. n = 6, error bars, mean ± SEM. P values were calculated using ordinary two-way ANOVA with Dunnett's multiple comparisons test.

(D) siRNA knockdown of RBM15, RBM15B, SPEN, and control in HEK293 cells followed by strand-specific RT-qPCR to measure viral cRNA. siRNA transfection was performed for 72 h followed by CedV infection (MOI 0.1) for indicated timepoints post-infection. Data is normalized to copy number at 0 hpi for each knockdown condition. n = 6, error bars, mean ± SD. P values were calculated using two way Anova with Dunnett's multiple comparisons test.

(E) siRNA knockdown of RBM15, RBM15B, SPEN, and control in HEK293 cells followed by strand-specific RT-qPCR to measure viral gRNA. siRNA transfection was performed for 72 h followed by CedV infection (MOI 0.1) for indicated timepoints post-infection. Data is normalized to copy number at 0 hpi for each knockdown condition. n = 6, error bars, mean ± SD. P values were calculated using two way Anova with Dunnett's multiple comparisons test.

**Supplementary Figure 5. m$^6$A modifications in CedV L mRNA are hypomethylated following SPEN depletion**

(A) Cell viability assay in STM2457-treated HEK293 cells. Data is normalized to DMSO vehicle. n = 6, error bars, mean ± SD.

(B) siRNA knockdown of YTHDF1, -2, or -3 in HEK293 cells followed by RT-qPCR to measure mRNA expression and evaluate knockdown efficiency. Data is normalized to TUBA1A using $\Delta\Delta C_t$ method and is relative to non-template control. n = 6, error bars, mean ± SD. P values were calculated using one-way Anova with Dunnett's multiple comparisons test.

(C) Barplot displaying percentages for human or CedV mapped reads from nanopore direct RNA sequencing of poly(A)+ RNA from SPEN knockdown or control HEK293 cells.

(D) Scatterplot displaying Pearson correlation of modification rates in CedV mRNA from biological replicates for control samples.

(E) Scatterplot displaying Pearson correlation of modification rates in CedV mRNA from biological replicates for SPEN knockdown samples.

Supplementary Data Figure 1

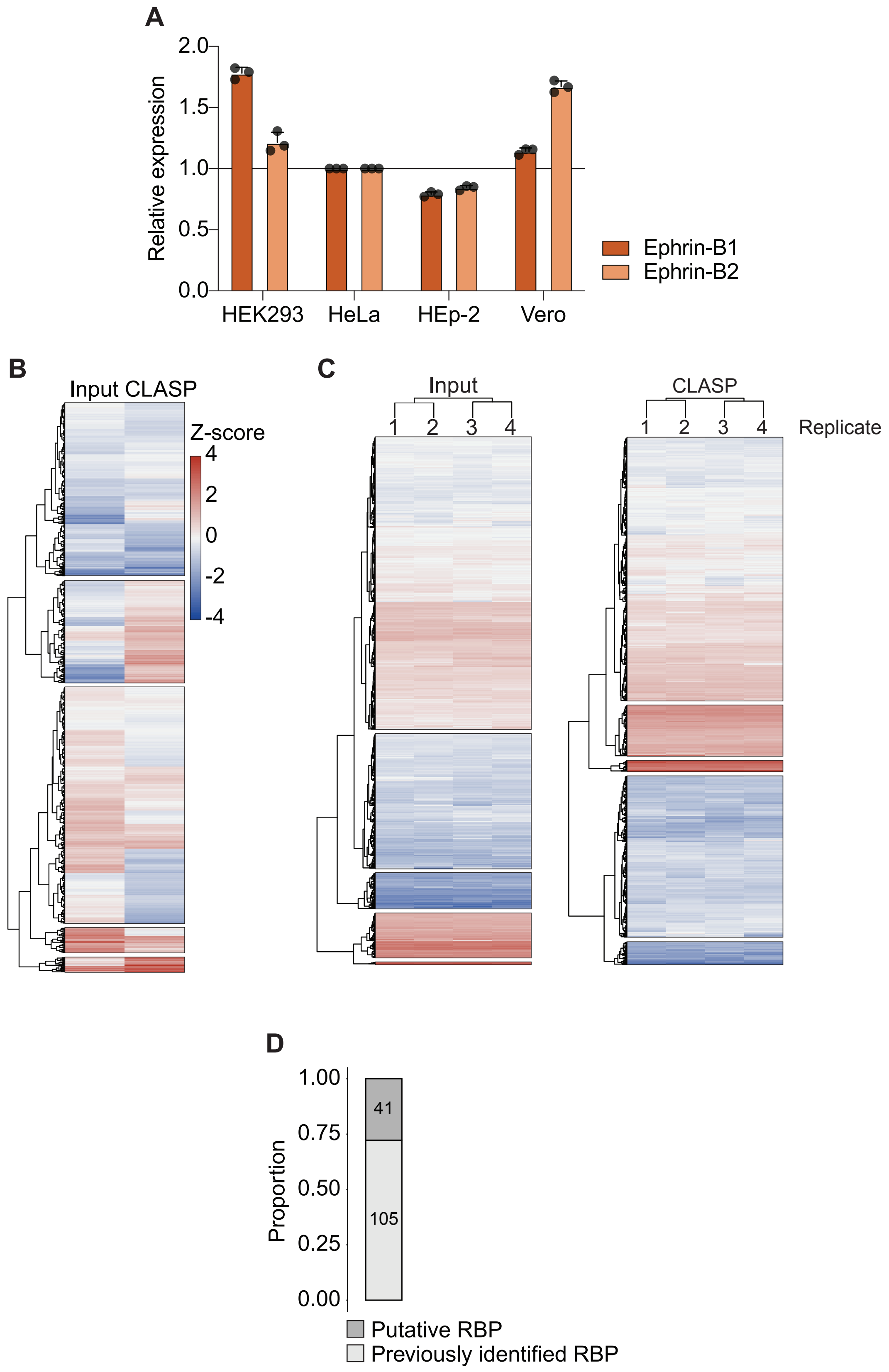

### Supplementary Data Figure 2

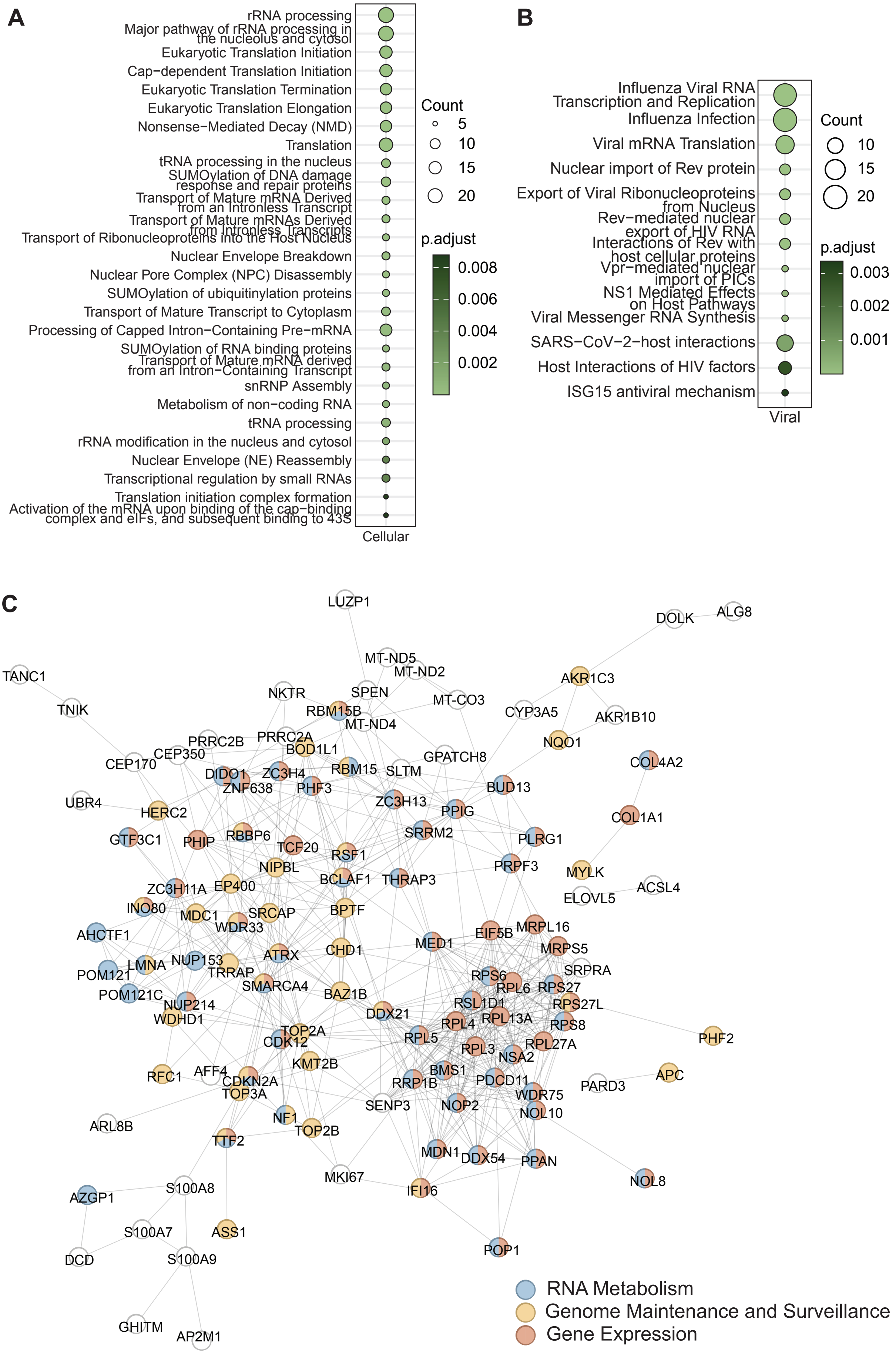

Supplementary Data Figure 3

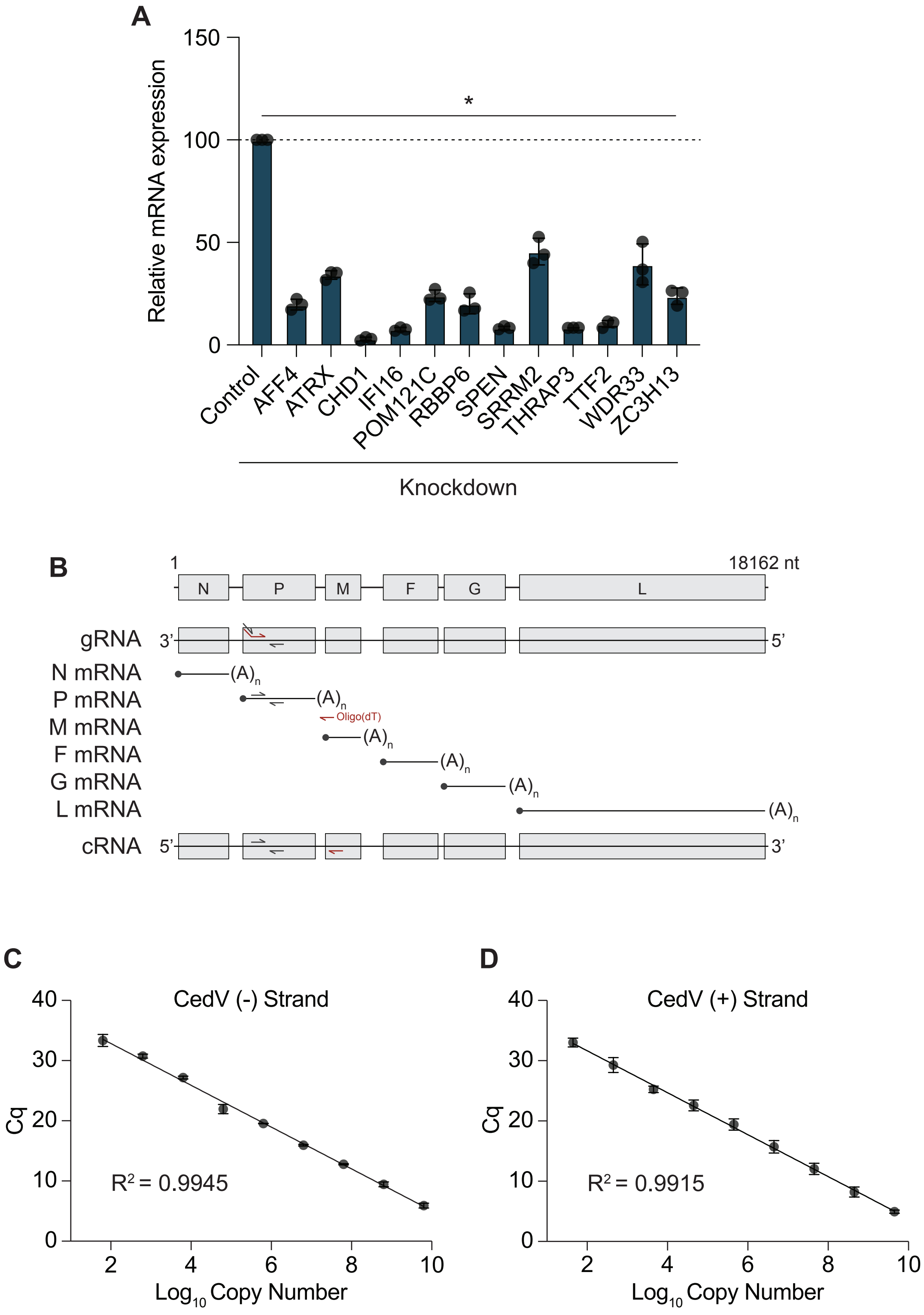

Supplementary Data Figure 4

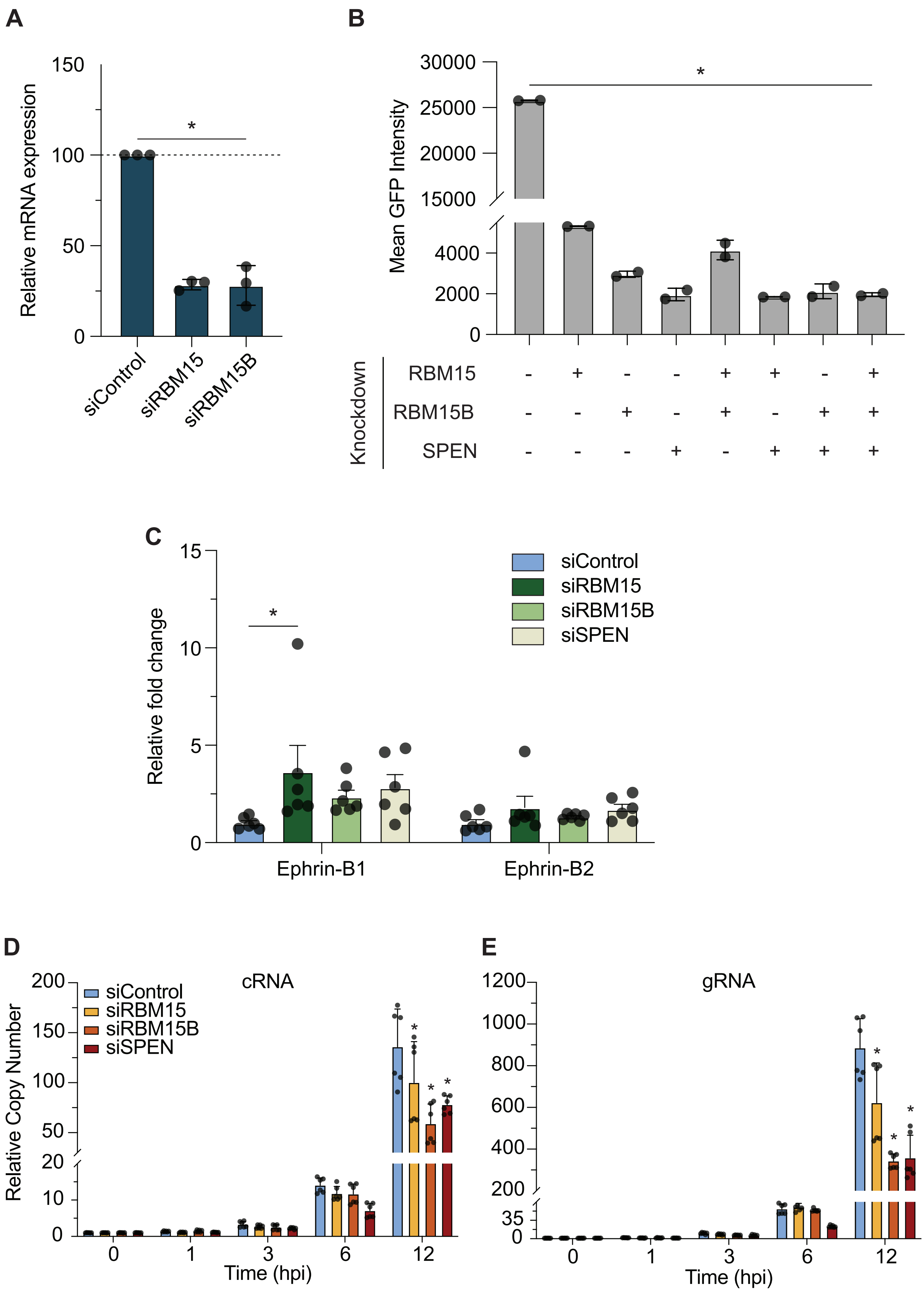

Supplementary Data Figure 5

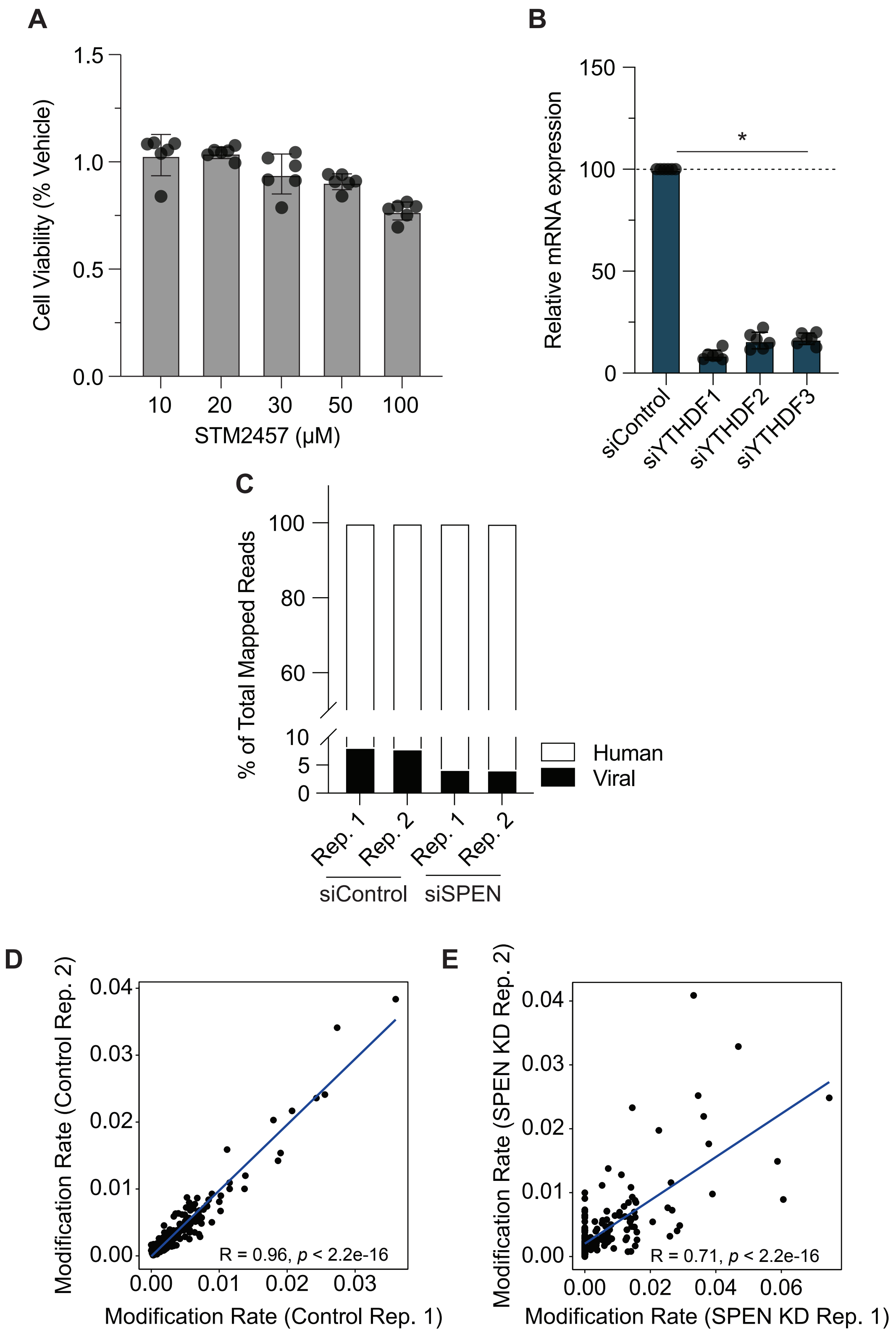
